# Detecting adaptive differentiation in structured populations with genomic data and common gardens

**DOI:** 10.1101/368506

**Authors:** Emily B. Josephs, Jeremy J. Berg, Jeffrey Ross-Ibarra, Graham Coop

## Abstract

Adaptation in quantitative traits often occurs through subtle shifts in allele frequencies at many loci, a process called polygenic adaptation. While a number of methods have been developed to detect polygenic adaptation in human populations, we lack clear strategies for doing so in many other systems. In particular, there is an opportunity to develop new methods that leverage datasets with genomic data and common garden trait measurements to systematically detect the quantitative traits important for adaptation. Here, we develop methods that do just this, using principal components of the relatedness matrix to detect excess divergence consistent with polygenic adaptation and using a conditional test to control for confounding effects due to population structure. We apply these methods to inbred maize lines from the USDA germplasm pool and maize landraces from Europe. Ultimately, these methods can be applied to additional domesticated and wild species to give us a broader picture of the specific traits that contribute to adaptation and the overall importance of polygenic adaptation in shaping quantitative trait variation.

## Introduction

Determining the traits involved in adaptation is crucial for understanding the maintenance of variation (Mitchell-Olds *et al.* 2007), the potential for organisms to adapt to climate change (Bay *et al.* 2017; Aitken *et al.* 2008), and the best strategies for breeding crops or livestock (Howden *et al.* 2007; Takeda and Matsuoka 2008). There are many examples of local adaptation from reciprocal transplant experiments (Hereford 2009; Leimu and Fischer 2008) that tell us about fitness in a specific environmental context, but these experiments are less informative about how past evolutionary forces have shaped present day variation (Savolainen *et al.* 2013). Instead, quantifying the role of adaptation in shaping current phenotypic variation will require comparing observed variation with expectations based on neutral models (Leinonen *et al.* 2008). With the growing number of large genomic and phenotypic common garden datasets, there is an opportunity to use these types of comparisons to systematically identify the traits that have diverged due to adaptation.

A common way of evaluating the role of spatially-variable selection in shaping genetic variation is to compare the proportion of the total quantitative trait variation among populations (*Q_ST_*) with that seen at neutral polymorphisms (*F_ST_*) (Spitze 1993; Prout and Barker 1993; Whitlock 2008). *Q_ST_ – F_ST_* methods have been successful at identifying local adaptation but have a few key limitations that are especially important for applications to large genomic and phenotypic datasets (Leinonen *et al.* 2013; Whitlock 2008). First, standard *Q_ST_ – F_ST_* assumes a model in which all populations are equally related (but see Whitlock and Gilbert 2012; Ovaskainen *et al.* 2011; Karhunen *et al.* 2013 for methods that incorporate different models of population structure). Second, rigorously estimating *Q_ST_* requires knowledge of the additive genetic variance *V_A_* both within and between populations (Whitlock 2008). Many studies skirt this demand by simply measuring the proportion of phenotypic variation partitioned between populations (“*P_ST_*”), either in natural habitats or in common gardens. However, replacing *Q_ST_* with *P_ST_* can lead to problems due to both environmental differences among natural populations and non-additive variation in common gardens (Pujol *et al.* 2008; Whitlock 2008; Brommer 2011). Third, *Q_ST_ –F_ST_* approaches are unable to evaluate selection in individuals or populations that have been genotyped but not phenotyped. In many cases it is more cost-effective to phenotype in a smaller panel and test for selection in a larger genotyped panel. Furthermore, there are a number of situations where it may be challenging to phenotype individuals of interest — for example, if individuals are heterozygous or outbred, cannot be easily maintained in controlled conditions, or are dead, they can be genotyped but not phenotyped. In these cases, the population genetic signature of adaptation in quantitative traits (“polygenic adaptation”) can be detected by looking for coordinated shifts in the allele frequencies at loci that affect the trait (Le Corre and Kremer 2012; Kremer and Le Corre 2012; Latta 1998).

Current approaches to detect polygenic adaptation in genomic data take advantage of patterns of variation at large numbers of loci identified in genome-wide association studies (GWAS) (Berg and Coop 2014; Turchin *et al.* 2012; Field *et al.* 2016). One approach, *Q_X_*, developed by Berg and Coop (2014), extends the intuition underlying classic *Q_ST_ – F_ST_* approaches by generating population-level polygenic scores — trait predictions generated from GWAS results and genomic data — and comparing these scores to a neutral expectation. However, methods for detecting polygenic adaptation using GWAS-identified loci are very sensitive to population structure in the GWAS panel (Berg and Coop 2014; Robinson *et al.* 2015; Novembre and Barton 2018; Berg *et al.* 2018; Sohail *et al.* 2018). Because GWAS in many systems are conducted in structured, species-wide panels (Atwell *et al.* 2010; Flint-Garcia *et al.* 2005; Wang *et al.* 2018), current methods for detecting polygenic adaptation are difficult to apply widely.

Here, we adapt methods for detecting polygenic adaptation to be used in structured GWAS panels and related populations. First, using a new strategy for estimating *V_A_*, we develop an extension of *Q_ST_ – F_ST_*, that we call *Q_PC_*, to test for evidence of adaptation in a heterogeneous, range-wide sample of individuals that have been genotyped and phenotyped in a common garden. We then develop an extension of *Q_X_* for use in structured GWAS populations where the panel used to test for selection shares population structure with the GWAS panel. We apply both of these methods to data from domesticated maize (*Zea mays* ssp. *mays*). Overall, we show that the method controls for false positive issues due to population structure and can detect selection on a number of traits in domesticated maize.

## Results

### Extending *Q_ST_ − F_ST_* to deal with complicated patterns of relatedness with *Q_PC_*

Our approach to detecting local adaptation is meant to ameliorate two main concerns of *Q_ST_ – F_ST_* analysis that limit its application to many datasets. First, many species-wide genomic datasets are collected from individuals that do not group naturally into populations, making it difficult to look for signatures of divergence between populations. Second, calculating *Q_ST_* requires an estimate of *V_A_*, usually done by phenotyping individuals from a crossing design.

We address these issues by using principal component analysis (PCA) to separate the kinship matrix, *K*, into a set of principal components (PCs) that can be used to estimate *V_A_* and an orthogonal set of PCs that can be used to test for selection. We base our use of PCA on the animal model, which is often used to partition phenotypic variance into the various genetic and environmental components among close relatives within populations (Henderson 1950, 1953; Thompson 2008). More generally, the animal model is a statement about the distribution of an additive phenotype if the loci contributing to the trait are drifting neutrally (see Ovaskainen *et al.* (2011); Berg and Coop (2014) for a recent discussion, and Hadfield and Nakagawa (2010) for a more general discussion of the relationship between the animal model and phylogenetic comparative methods).

We first use the animal model to describe how traits are expected to vary across individuals under drift alone. Let 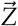 be a vector of measurements for a given trait across *M* individuals, taken in a common garden with shared environment. Assume for the moment that this trait is made up only of additive genetic effects, that environmental variation does not contribute to trait variation (*V_P_* = *V_A_*), and that measurements are without error (i.e. that 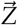 are breeding values). The animal model then states that 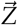 has a multivariate normal distribution:

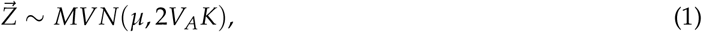

where *μ* is the mean phenotype, *V_A_* is the additive genetic variance, and *K* is a centered and standardized *M ×M* kinship matrix, where diagonal entries represent the inbreeding coefficients of individuals and off-diagonal cells represent the genotypic correlations between individuals (see Eq 16 in the methods). The kinship matrix describes how variation in a neutral additive genetic trait is structured among individuals due to variation in relatedness, while *V_A_* describes the scale of that variation.

Before discussing how we can use Eq. 1 to develop a test for adaptive divergence, we will show how this equation relates to *Q_ST_ – F_ST_*. If the individuals in our sample are grouped into a set of *P* distinct populations, then the kinship matrix can be used to generate an expectation of how trait variation is structured among populations under neutrality. To see this, consider that the vector of population mean breeding values 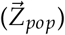 can be calculated from individual breeding values as 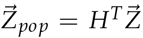, where the *p^th^* column of the *M × P* matrix *H* has entries of 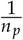 for individuals sampled from population *p*, and 0 otherwise (*n_p_* is the number of individuals sampled from population *p*). Because 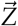 is multivariate normal, it follows that 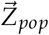 is as well, with

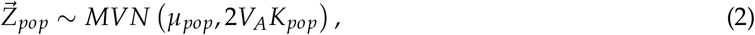

where *K_pop_* = *H^T^KH* and *μ_pop_* is the mean trait value across all populations.

Based on Eq. 2, we can calculate a simple summary statistic describing the deviation of 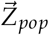 from the neutral expectation based on drift:

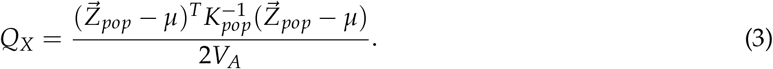

Under neutrality, *Q_X_* is expected to follow a *χ*^2^ distribution with *P –* 1 degrees of freedom (*μ* is not known *a priori* and must be estimated from the data, which expends a degree of freedom) (Berg and Coop 2014). If all *P* populations are equally diverged from one another, with no additional structure or inbreeding within groups, then *K_pop_* = *F_ST_ I*, where *F_ST_* is a measure of genetic differentiation between the populations and *I* is the identity matrix. Then, Eq. 3 simplifies to

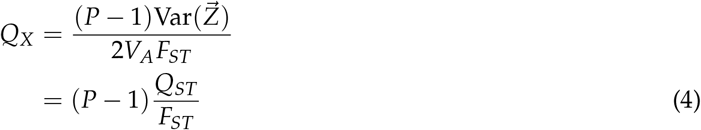

showing that the *Q_X_* statistic described in Eq 3 is the natural generalization of *Q_ST_ –F_ST_* to arbitrary population structure.

Here, we test for selection by looking for excess phenotypic divergence along the major axes of relatedness described by PCs instead of looking for excess divergence between populations, as is done in *Q_ST_ –F_ST_* and *Q_X_* analyses. For an example of how PCs relate to within and between population variation in a set of simulated populations, see Fig. 1. In these simulated populations, PCs 1 and 2 distinguish between-population variation for three populations while PC 3 separates out individuals within one population. While this is a simplified example compared to the populations analyzed later in the paper, it shows the intuition underlying our use of PCs to replace the within and between population structure used by *Q_ST_ − F_ST_*.

**Figure 1.**
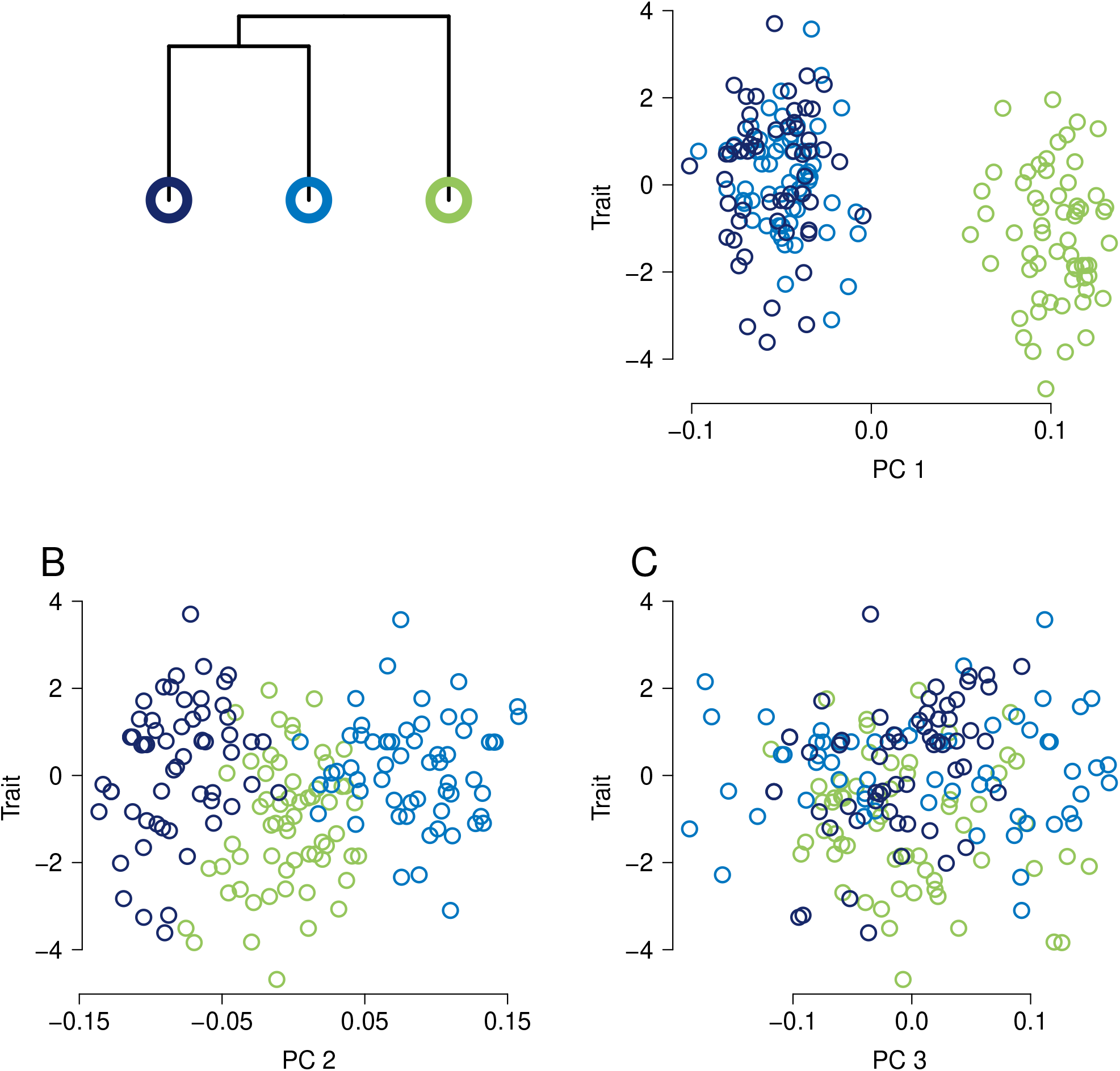
Relating PCs to within and between population variation. A conceptual figure based on three populations simulated following the dendrogram on left of plot. Each population has 70 individuals in it with some within-population variation in relatedness (not shown.). Panels A, B, and C, show the relationship between PCs 1, 2, and 3 and a neutrally evolving trait.

*Q_X_* can be linked to a PC based approach by noting that for any arbitrary *H* matrix (not just the type described above), *Q_X_* will follow a *χ*^2^ distribution and the degrees of freedom of this distribution will be equal to the number of linearly independent columns in *H*. We can generate a measure of excess divergence along specific PCs by replacing *H* with a matrix, *U*, of eigenvalues of the kinship matrix *K*. *U* is calculated from the eigendecomposition of *K*, such that, *K* = *U*Λ*U^T^* where 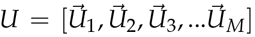 is the matrix of eigenvectors and Λ is a diagonal matrix with the eigenvalues of *K*. Here, we denote the *m^th^* eigenvector as 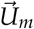 and the *m^th^* eigenvalue as *λ_m_* and we call our measure of excess divergence along PCs *Q_PC_*.

We will now walk through calculating *Q_PC_*. First we quantify the amount of divergence that occurs along PCs by projecting the trait described by 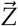 onto the eigenvectors of *K* by letting 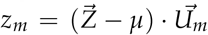. Intuitively, *z_m_* describes how much the traits 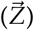 vary along the *m^th^* PC of the relatedness matrix *K*. *z_m_* can also be thought of as the slope of the relationship between 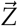 and the *m^th^* PC of *K*. Under a neutral model of drift (from Eq. 1) for each *m*:

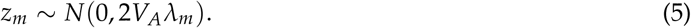

To compare *z_m_* across different PCs, we can calculate a standardized projection (*c_m_*):

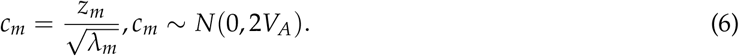

Crucially *c_m_* values represent deviations along linearly independent axes of neutral variation (PCs) and are independent from each other under neutrality; therefore, we can estimate *V_A_* using the variance of any set of *c_m_*. To develop a test analogous to *Q_ST_ − F_ST_*, we choose to declare projections onto the top 1: *R* of our eigenvectors 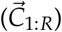 that explain broader patterns of relatedness to be “among population” axes of variation, and projections onto the lower *R* + 1: *M* of our eigenvectors (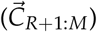) to be “within population” axes of variation (we will discuss our choice of *R* later).

Under neutrality, we expect that 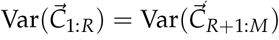. Adaptive differentiation among populations will increase the trait variance explained by the first PCs relative to the variance explained by later PCs and 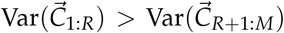. Note that 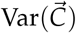 is the same as 𝔼 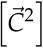 since the mean of 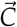 is 0 based on Eq. 6. We can test for differences between the variance of projections onto early PCs and the variance of projections later PCs using an *F* test:

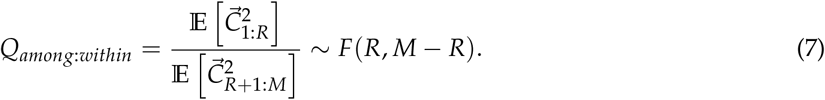

We focus on the upper tail of the F distribution, as we are interested in testing for evidence of selection contributing to trait divergence. Rejection of the null thus indicates excess trait variation in the first *R* PCs beyond an expectation based on the later *M – R* PCs. All together, this test allows us to detect adaptive trait divergence across a set of lines or individuals without having to group these individuals into specific populations.

We can also calculate variance along specific PCs and compare divergence along specific PCs to the additive variance estimated using the lower *R*: *M* eigenvectors. Looking at specific PCs will be useful for identifying the specific axes of relatedness variation that drive adaptive divergence as well as for visualizing results. For a given PC, *S*:

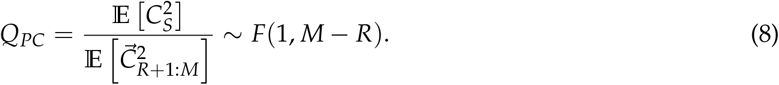

Again we test only in the upper tail of the distribution. The rejection of the null corresponds to excess variance along the *s^th^* PC beyond expectations based on variance along lower PCs. Eqs. 7 & 8 are valid for any values of S, R, and M as long as *R* > *S* and *M* > *R*. However, picking values of *S*, *R*, and *M* may not be trivial. In our subsequent application of this test, we choose to test for excess differentiation along the first set of PCs that cumulatively explain 30% of the total variation in relatedness. However, an alternative that we do not explore here would be selecting the set of PCs to use with methods from Bryc *et al.* (2013) or the Tracy-Widom distribution discussed in Patterson *et al.* (2006).

### Testing for selection with Q_PC_ in a maize mapping panel

We applied *Q_PC_* to test for selection in a panel of 240 inbred maize lines from the GWAS panel developed by Flint-Garcia *et al.* (2005). The GWAS panel includes inbred lines meant to represent the diversity of temperate and tropical lines used in public maize breeding programs, and these lines were recently sequenced as part of the maize HapMap 3 project Bukowski *et al.* (2017). In Fig. 2A we plot the relatedness of all maize lines on the first two PCs. The first PC explains 2.04% of the variance and separates out the tropical from the non-tropical lines, while the second PC explains 1.90% of the variance and differentiates the stiff-stalk samples from the rest of the dataset (stiff-stalk maize is one of the major heterotic groups used to make hybrids (Mikel and Dudley 2006)). While previous studies have used relatedness to assign lines to subpopulations, not all individuals can be easily assigned to a subpopulation and there is a fair amount of variation in relatedness within subpopulations (Flint-Garcia *et al.* 2005) (Fig. 2A), suggesting that using PCs to summarize relatedness will be useful for detecting adaptive divergence.

**Figure 2.**
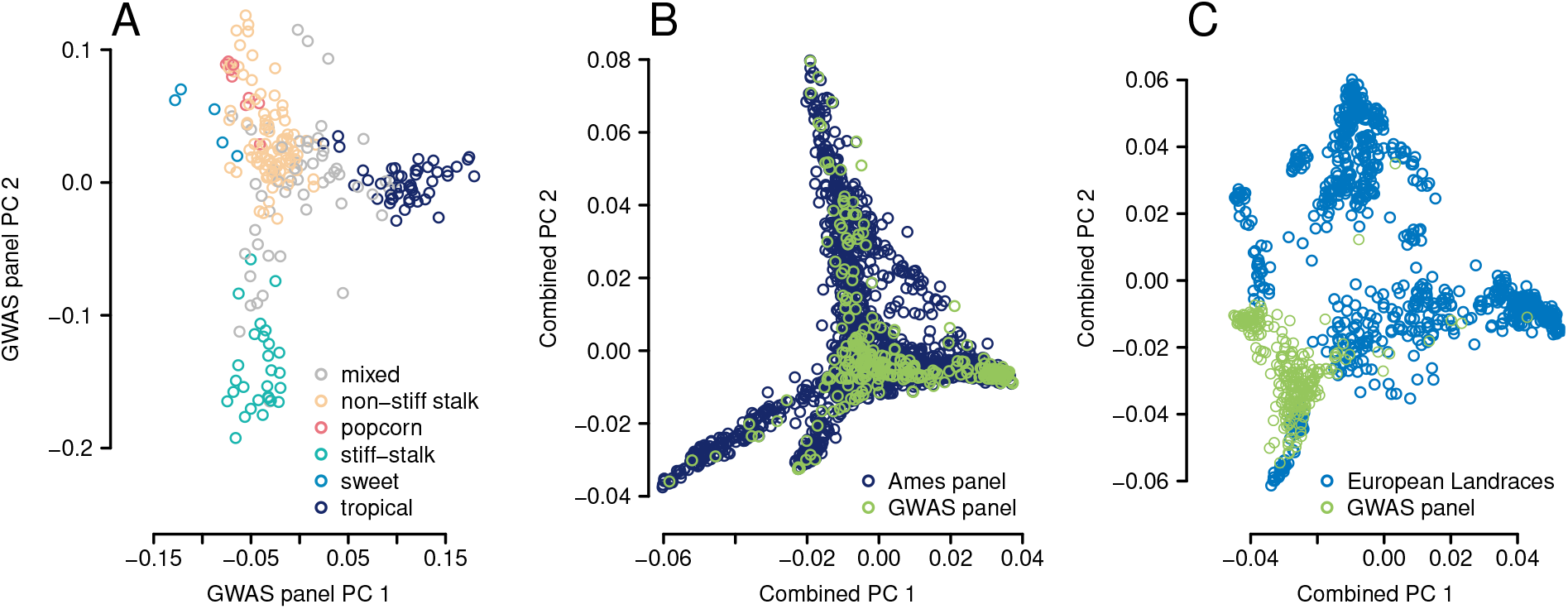
Structure in the maize populations. These plots show the first two principal components of population structure (the eigenvectors of the kinship matrix) for various maize panels included in this paper. A) 240 maize lines from the “GWAS panel” that were used in the trait *Q_PC_* analysis. Each point represents an inbred line and points are colored by their assignment to subpopulations from (Flint-Garcia *et al.* 2005). B) The GWAS panel from (A) along with the 2,704 inbred maize lines of the Ames diversity panel (Romay *et al.* 2013). C) The GWAS panel from (A) along with 906 European maize landraces from (*Unterseer et al. 2016)*

We first validated that *Q_PC_* would work on this panel by testing *Q_PC_* on 200 traits that we simulated under a multivariate normal model of drift based on the empirical kinship matrix, assuming *V_A_* = 1. As expected, from Eq. 6, the variance in the standardized projections onto PCs (*c_m_*) of these simulated traits centered on 1, and, across the 22 PCs tested in 200 simulations, only 200 tests (4.5%) were significant at the *p* < 0.05 level before correcting for multiple testing. Adding simulated environmental variation (*V_E_* = *V_A_*/10 and *V_E_* = *V_A_*/2) to trait measurements increased the variance of *c_m_*, with this excess variance falling disproportionately along the later PCs (those that explain less variation in relatedness). These results suggest that unaccounted *V_E_* increases estimated variance at later PCs, ultimately increasing the amount of variance along earlier PCs that will appear consistent with neutrality and reducing power to detect selection. However, this reduction in power can be minimized by controlling environmental noise — for example by measuring line replicates in a common garden or best unbiased linear predictions (BLUPs) from multiple environments (See Appendix 1 for a more extensive treatment of *V_E_*).

We tested for selection along 22 PCs for 22 traits that, themselves, are estimates of the breeding value (BLUPs) of these traits measured across multiple environments (Hung *et al.* 2012). These 22 traits include a number of traits thought to be important for adaptation to domestication and/or temperate environments in maize, such as flowering time (Swarts *et al.* 2017), upper leaf angle (Duvick 2005), and plant height (Peiffer *et al.* 2014; Duvick 2005). After controlling for multiple testing using an FDR of 0.05, we found evidence of adaptive divergence for three traits: days to silk, days to anthesis, and node number below ear (Fig. 3A). These results are generally robust to the choice of PCs used for the denominator of Eq. 8 used to estimate *V_A_* (Fig. S3). We plot the relationship between PC1 and two example traits to illustrate the data underlying these signals of selection. In Fig. 3B, we show a relationship between PC1 and Kernel Number that is consistent with neutral processes and in Fig. 3C we show a relationship between PC 1 and days to silk that is stronger than would be expected due to neutral processes and is instead consistent with diversifying selection. We detected evidence of diversifying selection on various traits along PC1, PC2, and PC10. While PC 1 and PC2 differentiate between known maize subpopulations (Fig. 2A), PC 10 separates out individuals within the tropical subpopulation, so our results are consistent with adaptive divergence contributing to trait variation within the tropical subpopulation (Fig. S2).

**Figure 3.**
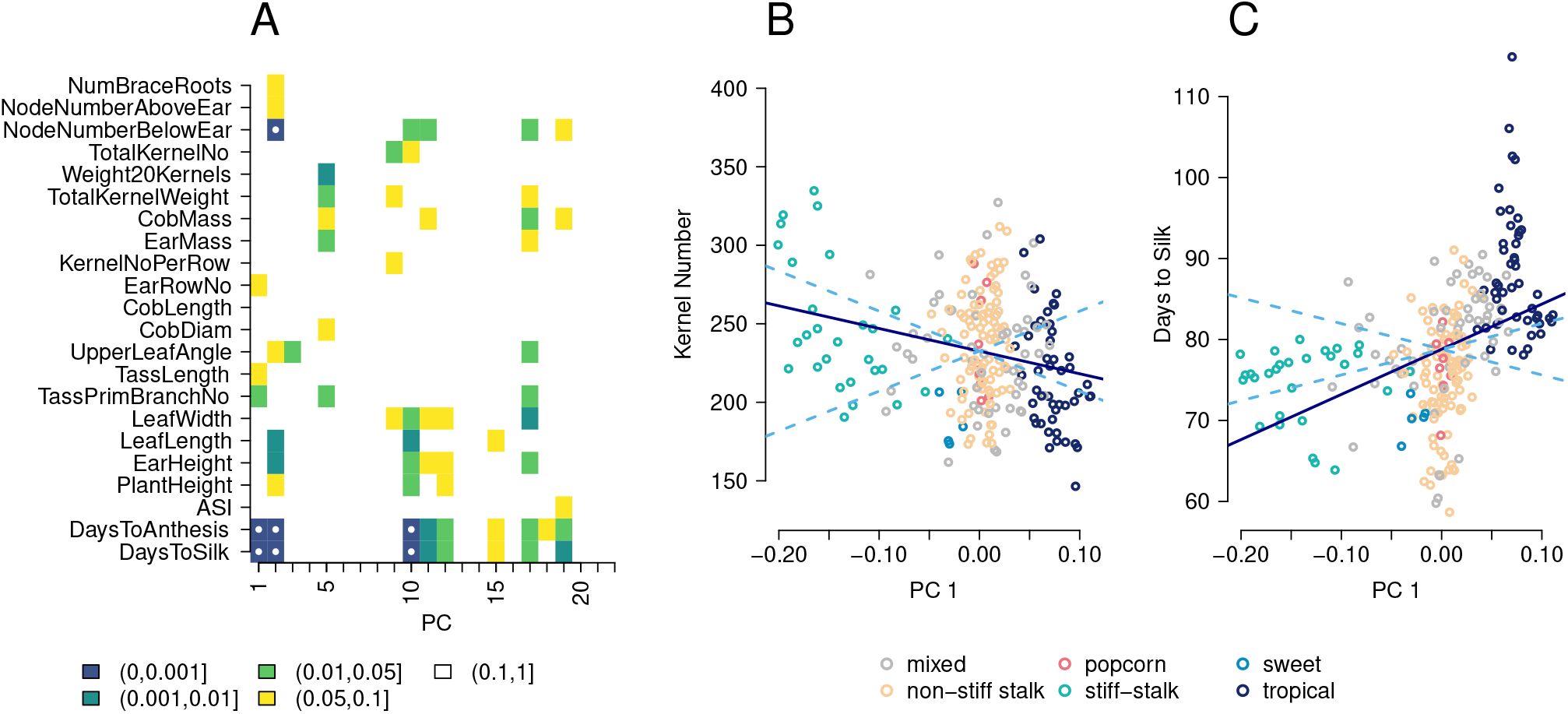
Detecting adaptation within the GWAS panel with *Q_PC_*. A) A heatmap showing results from *Q_PC_* on the first 22 PCs (x-axis) for 22 traits (y-axis). Squares are colored by their p value. If the false discovery rate (’q value’) corresponding to that p value is < 0.05 there is a white dot in the square. B) Total kernel number per cob plotted against the first principal component of relatedness (PC 1). Each point represents a line in the GWAS panel, colored by its membership in a subpopulation (same colors as Fig. 2A). The solid line shows the linear regression of the trait on PC 1 and the dashed lines show the 95% confidence interval of linear regressions expected under neutrality. Note that the linear regression is not the same as the F test done in *Q_PC_*, and that we plot these lines for visualization purposes only. C) Similar to B, but showing days to silk on the Y axis. The regression line for days to silk against PC 1 (solid line) falls outside the expectation based on neutrality (dotted line), consistent with selection increasing trait divergence across PC 1.

### Detecting selection in un-phenotyped individuals using polygenic scores

Extending the method described above to detect selection in individuals or lines that have been genotyped but not phenotyped will allow the study of polygenic adaptation when phenotyping is expensive or impossible. Here we outline methods for detecting selection in individuals that have been genotyped but not phenotyped (we refer to these individuals as the“genotyping panel”). We build on methods developed in Berg and Coop (2014) and Berg *et al.* (2017) and extend them to test for adaptive divergence along specific PCs and in the presence of population structure shared between the GWAS panel and the genotyping panel. The main difference between this test and *Q_PC_* as described previously is that we now test for selection on polygenic scores, not on measured phenotypes. Specifically, if we have a set of *n* independent, trait-associated loci found in a GWAS, we can write the polygenic score for individual or line *i* as:

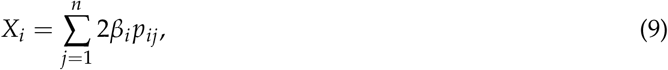

where *β_j_* is the additive effect of having an alternate allele of the *j^th^* locus, and *p_ij_* is the alternate allele’s frequency within the *i^th^* individual or line (i.e., half the number of allele copies in a diploid individual).

However, when there is shared population structure between the GWAS panel and the genotyping panel, there are two concerns about testing for selection on polygenic scores made for the genotyping panel:

1. If we have already found a signal of selection on our phenotypes of interest in the GWAS panel, then a significant test could simply reflect this same signal and not independent adaptation in the genotyping panel.
2. Controls for population structure in the GWAS could bias the loci identified and the effect sizes estimated for these loci, leading to false positive signals of selection in the genotyping panel.

This second point is worth considering carefully. Modern GWAS control for false-positive associations due to population structure, often by incorporating a random effect based on the kinship matrix into the GWAS model (Yu *et al.* 2006). However, controlling for population structure will bias GWAS towards finding associations at alleles whose distributions do not follow neutral population structure and towards missing true associations with loci whose distributions do follow population structure (Atwell *et al.* 2010). Because of this bias, the loci detected may not appear to have neutral distributions in the GWAS panel or, crucially, in any additional set of populations that share structure with the GWAS panel.

We illustrate this problem by looking at the relationship between PC 1 and polygenic scores for a trait that has been simulated to be evolving neutrally in the GWAS panel and a set of related maize lines (Fig. 4A). While there is no strong relationship between the simulated trait and PC 1 (Fig. 4A), the polygenic scores calculated from GWAS results do show a strong correlation with PC1 (Fig. 4B). Note that the simulated trait values and polygenic scores have been standardized by dividing by the square root of their respective *V_A_*s estimated from the lower PCs, so this erroneous correlation not simply a result of the inflation of effect sizes in the GWAS due to false positives and the winner’s curse but represents an issue caused by shared population structure between the GWAS panel and the genotyping panel. While these plots show data for 1 simulation, the magnitude of the slope of the relationship between standardized polygenic score and PC 1 was greater than the magnitude of the slope of the relationship between standardized simulated trait and PC 1 for 162 out of 200 simulations.

**Figure 4.**
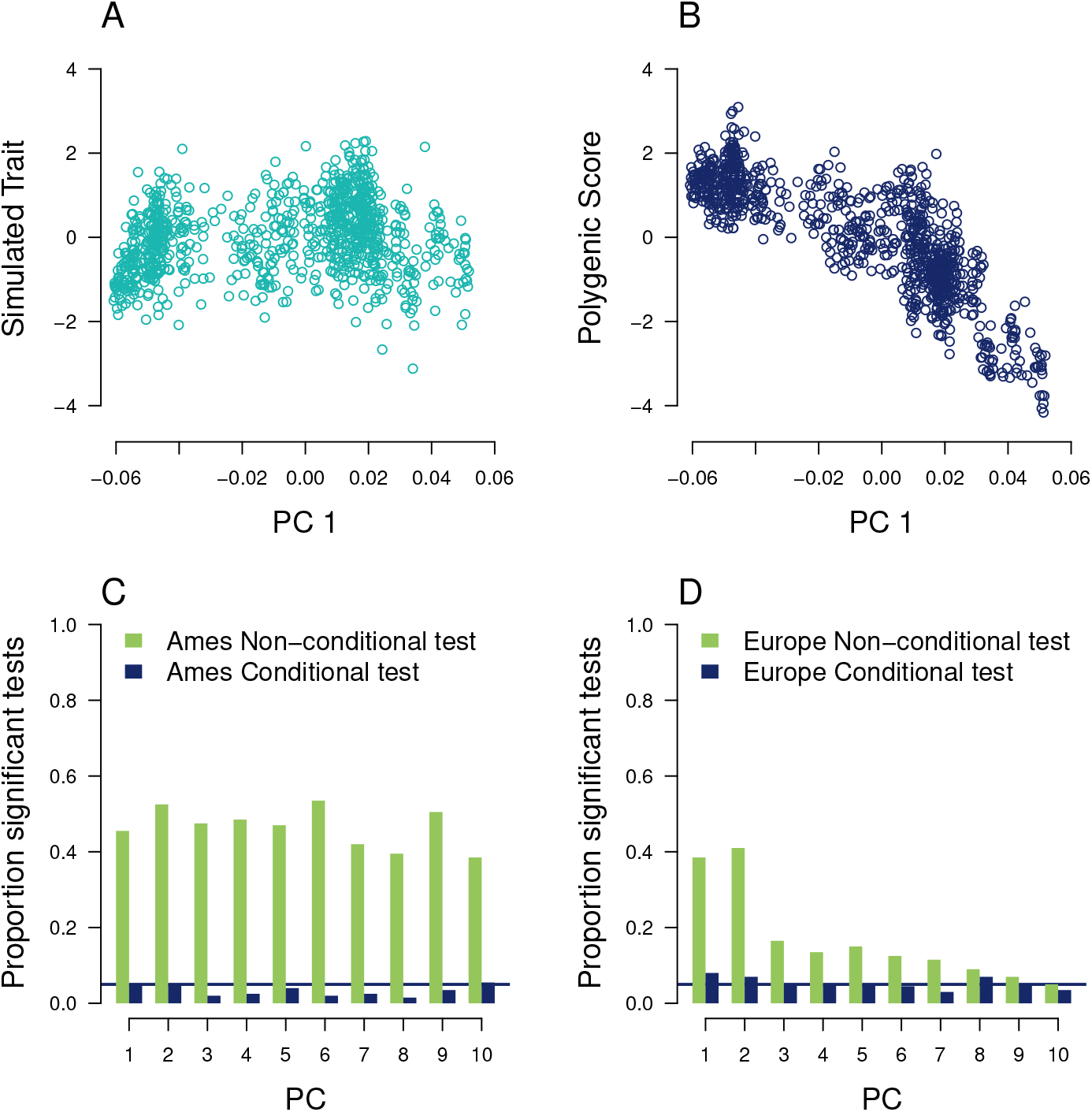
Simulations of *Q_PC_* on polygenic scores. A) The relationship between a neutrally simulated trait and PC 1 the European landraces. Each point represents one landrace and the trait value has been divided by 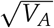 so that it can be compared to traits with different values of *V_A_* B) The relationship between polygenic scores calculated for the same trait as in (A), based on a GWAS in the GWAS panel. This relationship between polygenic scores and PC is what is used in the non-conditional *Q_PC_* test. As in (A), each point represents one maize individual and trait has been standardized by *V_A_*. C) The proportion of 200 neutral simulations that were significant at the *p* < 0.05 level for the non-conditional *Q_PC_* test and the conditional *Q_PC_* test. A horizontal line is plotted at 0.05, to show the proportion of significant tests expected under the null hypothesis D) The same information, this time for the European landraces.

Here, we control for shared structure between the GWAS and genotyping panels by conditioning on the estimated polygenic scores in the GWAS panel 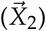 when assessing patterns of selection on the polygenic scores of a genotyping panel 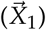. Specifically, following the multivariate normality assumption (Eq. 1), we model the combined vector of polygenic scores in both panels as:

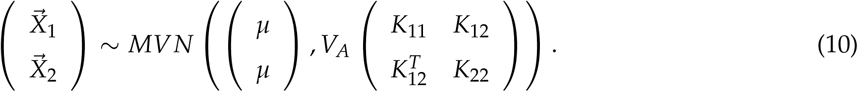

where, *μ* is the mean of the combined vector [*X*_1_, *X*_2_], *K*_11_ and *K*_22_ are the kinship matrices of the genotyping and GWAS panels, and *K*_12_ is the set of relatedness coefficients between lines in the genotyping panel (rows) and GWAS panel (columns). Note that the combination of the four kinship matrices in the variance term of Eq. 10 is equivalent to the kinship matrix of all individuals in the genotyping and GWAS panels. We discuss the mean centering of these matrices in Appendix 3.

The conditional multivariate null model for polygenic scores in the genotyping panel conditional on the GWAS panel is then

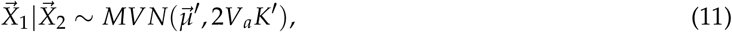

Where 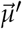 is a vector of conditional means with an entry for each sample in the genotyping panel,

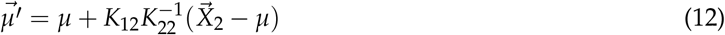

and *K′* is the relatedness matrix for the genotyping panel conditional on the matrix of the GWAS panel,

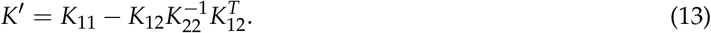

Following Eq. 6 and Eq. 8 we can test for excess variation along the PCs of *K′*, using the difference between polygenic scores 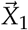 and the conditional means 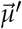 as the phenotype. Specifically, if 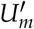 and 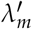 are the *m^th^* eigenvector and eigenvalue of *K′*, then

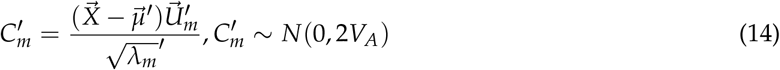

and

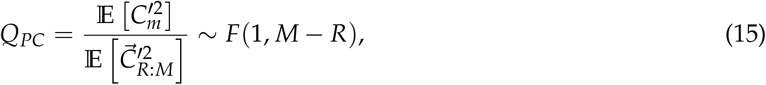

where *R* > *m* and *M* > *R*. We will refer to the conditional version of the test as ‘conditional *Q_PC_*’.

It is worth taking some time to discuss how the conditional test controls for the two issues due to shared structure that discussed previously. First, by incorporating the polygenic scores of individuals in the GWAS panel into the null distribution of conditional *Q_PC_*, we are able to test directly for adaptive divergence that occurred in the genotyping panel. Berg *et al.* (2017) also uses the conditional test in this manner. Second, the conditional test forces the polygenic scores of individuals in the genotyping panel into the same multivariate normal distribution as the polygenic scores of individuals in the GWAS panel. Since the polygenic scores of GWAS individuals will include the ascertainment biases expected due to controls for structure in the GWAS, these biases will be incorporated into the null distribution of polygenic scores expected under drift. Now, we will only detect selection if trait divergence exceeds neutral expectations based on this combined multivariate normal distribution.

### Applying Q_PC_ to polygenic scores in North American inbred maize lines and European landraces

First, we conducted a set of neutral simulations to assess the ability of the conditional *Q_PC_* test to control for false positives due to shared structure. We applied both the conditional and original (“non-conditional”) *Q_PC_* test to detect selection on polygenic scores constructed from simulated neutral loci in two panels of maize genotypes that have not been extensively phenotyped: a set of 2,815 inbred lines from the USDA that we refer to as ‘the Ames panel’ (Romay *et al.* 2013) and a set of 906 individuals from 38 European landraces (Unterseer *et al.* 2016). We chose these two panels to evaluate the potential of conditional *Q_PC_* to control for shared population structure when the problem is severe, as in the Ames panel (Fig. 2B), and moderate, as in the European landraces (Fig. 2C). In addition, we expect that the evolution of many quantitative traits has been important for European landraces as they adapted to new European environments in the last ~500 years (Unterseer *et al.* 2016; Tenaillon and Charcosset 2011).

False positive signatures of selection were common when using the original *Q_PC_* based on relatedness within the genotyping panel to test for selection on polygenic scores based on loci simulated under neutral processes (Fig. 4C, D, Fig. S4) These false positive signals were common even though the GWAS used to identify the loci used to make polygenic scores both had low power to detect associations and had high false positive rates. (Fig. S5). The increase in false positives due to shared structure persisted to much later PCs in the Ames panel than in the European landraces, likely because the extent of shared structure is more pervasive for the Ames panel. However, the conditional *Q_PC_* test appeared to control for false positives in both the Ames panel and the European landraces (Fig. 4A, B.).

We then conducted GWAS on 22 traits in the GWAS panel. We used a p value cutoff of 0.005 to choose loci for constructing polygenic scores. This cutoff is less stringent then the cutoffs standardly used in maize GWAS (Peiffer *et al.* 2014; Romay *et al.* 2013), but allowed us to detect a large number of loci that we could use to construct polygenic scores. After thinning the loci for linkage disequilibrium, we found associations for all traits with an average of 350 associated SNPs per trait (range 254 – 493, supp figures). We used these SNPs to construct polygenic scores for lines in the Ames panel and individuals in the European landraces following Eq 9.

When we applied the original (non-conditional) test from Eq. 8 to detect selection in the Ames panel on 22 traits for 182 PCs (4,004 tests total), we uncovered signals of widespread polygenic adaptation (Fig. S6A, Fig. S7A). In contrast, conditional *Q_PC_* on the same set of traits and PCs found no signatures of polygenic adaptation in the Ames panel that survived control for multiple testing (Fig S6B, Fig. S7B). The lack of results in the conditional test is unsurprising because the GWAS panel’s population structure almost completely overlaps the Ames panel (Fig. 2B), so once variation in the GWAS panel is accounted for in the conditional test, there is likely little differentiation in polygenic scores left to test for selection. We report these results to highlight the caution that researchers should use when applying methods for detecting polygenic adaptation to genotyping panels that share population structure with GWAS panels.

In the European landraces, we conducted *Q_PC_* on 22 traits across 17 PCs (374 tests total) and while we detected selection on a number of traits, as with the Ames panel, none of these signals were robust to controlling for multiple testing using a false-discovery rate approach (Fig. S7D). However, we report the results that were significant at an uncorrected level in Fig. 5A to demonstrate how these types of selective signals could be visualized with these approaches. In Fig. 5B, we show the relationship between conditional PC1 (*U*_1_) and the difference between polygenic score for the number of brace roots and a conditional expectation 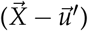 which was our strongest signal of selection in the panel.

**Figure 5.**
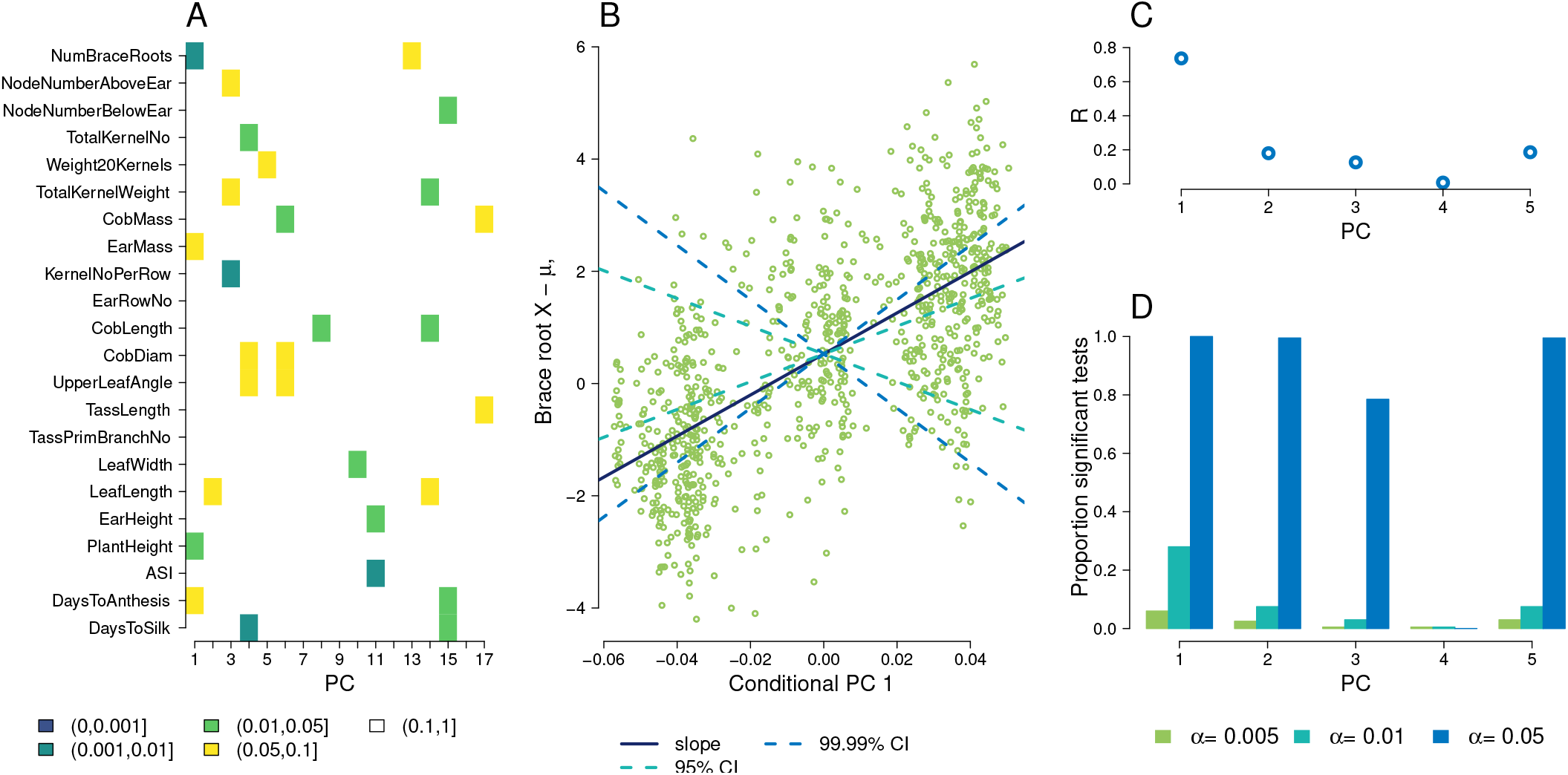
Detecting adaptation within the European landraces with *Q_PC_* on polygenic scores. A) A heatmap showing results from the conditional *Q_PC_* on the first 17 PCs (x-axis) for 22 traits (y-axis). Squares are colored by uncorrected p value if p<0.1. B) Marginal evidence of selection on brace root number. The difference between polygenic score for brace root and the conditional mean 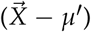 is plotted against the conditional PC 1 for each line. The solid line shows the observed linear relationship between conditional PC 1 and 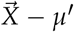 and the dotted lines show the 95% confidence interval for this slope based on neutral expectations uncorrected for multiple testing and the 99.99% confidence interval corresponding to a Bonferonni correction for multiple testing. C) The absolute value of the correlation coefficient (R) between PCs and latitude. D) The proportion of significant tests for traits simulated under three different selection strengths along latitude.

We conducted power simulations by shifting allele frequencies of GWAS-identified loci along a latitudinal selective gradient in the European landraces (see Methods section for details). When selection was strong (selection gradient *α* = 0.05), we detected signals of selection in all 200 simulations along the first conditional PC, which had the strongest association with latitude. When selection was moderate (*α* = 0.01) we detected selection in 57 of 200 simulations (Fig. 5C,D). These results suggest that there is power to detect selection on polygenic scores with *Q_PC_* in the European landraces if selection actually occurs on the loci used to make these polygenic scores.

## Discussion

In this paper we have laid out a set of approaches that can be used to study adaptation and divergent selection using genomic and phenotypic data from structured populations. We first described a method, *Q_PC_*, that can be used to detect adaptive trait divergence in a species-wide sample of individuals or lines that have been phenotyped in common garden and genotyped. We demonstrated this method using a panel of phenotyped domesticated maize lines, showing evidence of selection on flowering time and node number below ear. Second, we present an extension of *Q_PC_* that can be applied to individuals related to the GWAS panel that have not themselves been phenotyped using a conditional test to avoid confounding due to shared population structure. We showed that this test is robust to false-positives due to population structure shared between the GWAS panel and the genotyping panel and that it has power to detect selection. We applied this method to two panels of maize lines and showed marginal evidence of selection on a number of traits, but these signals were not robust to multiple testing corrections. Overall, the methods described and demonstrated here will be useful to a wide range of study systems.

While we were able to use *Q_PC_* to detect diversifying selection on phenotypes in the GWAS panel of 240 inbred lines, we were unable to detect similar patterns using polygenic scores for the Ames panel and European landraces (after controlling for multiple testing). This lack of selective signal was expected in Ames because the high overlap in relatedness between the Ames panel and the GWAS panel reduces power to detect selection in the Ames panel alone. However, our simulations showed that we did have power to detect moderate to strong selection acting on GWAS-associated loci in European landraces, and we expect that adaptation to European environments has contributed to trait diversification (Unterseer *et al.* 2016). There are a few factors that could explain our inability to detect selection on polygenic scores for European landraces. First, the polygenic scores we constructed used GWAS results from traits measured in North American environments. If there is *G ×E* for these traits, we may not be measuring traits that are actually under selection in Europe. Second, it is likely that in our small GWAS panel (n = 263 or 281) we are underpowered to detect most causal loci and so our predictions are too inaccurate to pull out a signal of selection. All together, our results suggest that while GWAS are undoubtedly useful to identify loci underlying traits, an analysis of phenotypes expressed in a common environment will often be the most powerful approach for detecting adaptation, especially in systems with under-powered GWAS.

Our proposed methods use principal component analysis (PCA) to separate out independent axes of population structure. There is a clear connection between PCA and average pairwise coalescent times (McVean 2009) and, because of this connection, PCA has been useful in a range of population genetic applications, including the detection and visualization of population structure (Patterson *et al.* 2006; Novembre *et al.* 2008), understanding the roles of population history and geography (Menozzi *et al.* 1978; Novembre and Stephens 2008), controlling for population structure in genome-wide association studies (Price *et al.* 2006). Here we use PCs to define broad axes of variation analogous to the between-population variation described in standard *Q_ST_ – F_ST_* tests while using later PCs to describe variation within populations and. In our application to maize data, we use arbitrary cutoffs to define which PCs represent between- and within-population variation. In the future, developing a rigorous method for choosing appropriate PCs will be useful, but for now, we suggest that researchers use their own knowledge of their study system and datasets to choose these PCs. We also caution that the presence of many close relatives in the sample could affect PCA, such that early PCs could end up representing variation within a large family and we suggest that researchers remove close relatives when using these methods. In addition, while PCs provide a useful way of separating signals, in some cases the constraints of PCA make the PCs unintuitive in terms of geography and environmental variables (Novembre and Stephens 2008). Therefore, it will also be useful to explore approaches like that outlined in Eq. 9 of Berg *et al.* (2018) that project trait values or polygenic scores onto environmental variables to test if the relationship between trait and environment is larger than would be expected due to drift.

There are a number of connections between the methods presented here and previous approaches.Ovaskainen *et al.* (2011); Karhunen *et al.* (2013) calculated a *Q_ST_ – F_ST_*-like measure of diversifying selection using the kinship matrix to model variation in relatedness among subpopulations. Their approach, however, is still reliant on identifying sub-populations and on using trait measurements in families or crosses to obtain estimates of *V_A_*. For single loci, a number of *F_ST_*-like approaches have been developed that use PCs to replace subpopulation structure to detecting individual outlier loci that deviate from a neutral model of population structure (Duforet-Frebourg *et al.* 2015; Luu *et al.* 2016; Galinsky *et al.* 2016; Chen *et al.* 2016). Our methods can be viewed as a a phenotypic equivalent to these locus-level approaches. In addition, Liu *et al.* (2018) have recently explored a related approach using projections of polygenic scores along PCs. Finally it may be useful to recast our method in terms of the animal model by splitting the kinship matrix into a ‘between population’ matrix described by early PCs and a ‘within population’ matrix described by later PCs. We could then detect selection by comparing estimates of *V_A_* for these two matrices. Such an animal-model approach may also offer a way to incorporate environmental variance in systems where replicates of the same genotype are not possible.

There are a number of caveats for applying *Q_PC_* to traits on additional systems and datasets that stem from issues in accurately estimating *V_A_* and it is important to carefully consider the assumption underlying *Q_PC_* that all traits are made up of additive combinations of allelic effects. If environmental variation contributes to trait variation, it will reduce the power of *Q_PC_* to detect diversifying selection because environmental variation will contribute most to variation at later PCs (Appendix 1). Additionally, additive-by-additive epistasis has the potential to contribute to false-positive signals of adaptation because additive-by-additive epistatic variation will contribute most to phenotypic variance along earlier PCs (Appendix 2). In general, non-additive interactions between alleles may cause difficulty for *Q_PC_* in systems, like maize, where traits are measured on inbred lines but selection occurs on outbred individuals. However, there is evidence that additive-by-additive variance will often be small compared *V_A_* within populations (Hill *et al.* 2008); for example, the genetic basis of flowering time variation in maize is largely additive (Buckler *et al.* 2009), suggesting that our conclusions about adaptive divergence in flowering time are likely robust to concerns about epistasis.

Our results highlight a number of issues with polygenic adaptation tests that depend on polygenic scores using GWAS-associated loci. As has been recently highlighted by Berg *et al.* (2018) and Sohail *et al.* (2018), structure in a GWAS panel can contribute to false signals of polygenic adaptation in polygenic scores constructed from the results of that panel. We observed that this problem is especially strong when there is shared population structure between the GWAS panel and the genotyping panel used to construct polygenic scores but that the use of a conditional test that accounts for shared structure between the two datasets can control for these false positives. There is potential for these methods to be used to address problems due to structure in GWAS panels in both non-human and human systems, although the conditional test approach would need to be adapted to the very large sample sizes used in human GWAS.

All together, the methods presented here provide an approach to detecting the role of diversifying selection in shaping patterns of trait variation across a number of species and traits. A number of further avenues exist for extending these methods. First, we applied this test to traits independently, but extending *Q_PC_* to incorporate multiple correlated traits will likely improve power to detect selection by reducing the number of tests done. In addition, this extension could allow the detection of adaptive changes in trait correlations. Second, these methods could be extended to take advantage of more sophisticated methods of genomic prediction than the additive model presented here (as in Beissinger *et al.* (2018); Liu *et al.* (2018)). Pursuing this goal will require carefully addressing issues related to linkage disequilibrium between marker loci. Overall, developing and applying methods for detecting polygenic adaptation in a wide range of species will be crucial for understanding the broad contribution of adaptation to phenotypic divergence.

## Materials and Methods

Analyses were done in R and we used the dplyr package (R Core Team 2018; Wickham *et al.* 2017). All code is available at https://github.com/emjosephs/qpc-maize.

### The germplasm used in this study

We analyzed three different maize diversity panels.

- **The GWAS panel**: The Major Goodman GWAS panel, also sometimes referred to as ‘the 282’ or ‘the Flint Garcia GWAS Panel’, contains 302 inbred lines meant to represent the genetic diversity of public maize-breeding programs (Flint-Garcia *et al.* 2005). Genotype-by-sequencing (GBS) data is available for 281 of these lines from Romay *et al.* (2013)and 7X genomic sequence from 271 of these lines is available from Bukowski *et al.* (2017). In addition, these lines have been phenotyped for 22 traits in multiple common garden experiments (Hung *et al.* 2012).
- **The Ames panel:** A panel of 2,815 inbred lines from the USDA that have been genotyped with GBS (Romay *et al.* 2013) at 717,588 SNPs.
- **The European landraces:** A panel of 906 individuals from 38 European landraces (31 Flint-type and 7 Dent-type) that were genotyped at 547,412 SNPs using an array (Unterseer *et al.* 2016).

### Q_PC_ **in the GWAS panel**

We tested for selection on 22 traits phenotyped in the GWAS panel. Best unbiased linear predictions (BLUPs) for these traits were sourced from Hung *et al.* (2012) and genomic sequence data from Bukowski *et al.* (2017). Out of the 302 individuals in the GWAS panel, we retained 240 individuals that had data for all 22 traits of interest and had genotype calls for >70% of the SNPs in the genomic dataset.

To construct a kinship matrix for the GWAS panel, we randomly sampled 50,000 SNPs from across the genome after removing sites that were missing any data or had unrealistic levels of heterozygosity (the proportion of heterozygous individuals exceeded 0.5). The allele frequencies (0, 0.5, or 1) for individuals at these 50,000 SNPs were arranged in an *MxN* matrix (referred to here as *G*) where *M* is the number of individuals (240) and *N* is the number of loci (50,000) such that *g_in_* is the entry on the *i^th^* row and *n^th^* column of *G* and it represents the genotype of the *i^th^* individual at the *n^th^* locus.

Then we centered the matrix using a centering matrix, *T*, which is an *M −* 1 by *M* matrix with 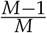 on the diagonal and 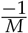 at all other cells. The matrix *TG* will be the matrix *G* that has been mean centered, such that if (*tg*)_*in*_is the entry for the *i^th^* individual and *n^th^* locus and *g*̄*n* is the mean allele frequency of the *n^th^* locus, (*tg*)_*in*_= *g_in_ – g*̄*n* Note that multiplying *G* by *T* also drops one individual from the kinship matrix to reflect the fact that by mean centering, we have lost one degree of freedom. We also standardized *G* by dividing by the square root of the expected heterozygosity of all loci, calculated by taking the mean of ϵ (1– ϵ) across all loci, where ϵ is the mean allele frequency of a locus. All together, we calculated *K* as the covariance of the centered and standardized matrix:

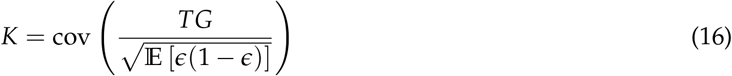

Each cell of *K* contains the covariance of genotypes between individuals, such that if *K_ij_* is the value of the *i^th^* row and *j^th^* column of *K* it represents the covariance between the *i^th^* and *j^th^* indviduals, such that,

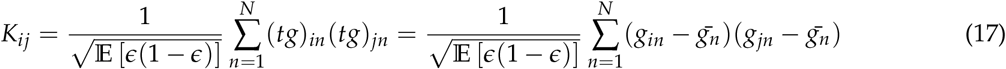

The principal components of the genotype matrix *G* were calculated from the eigenvectors of *K*. However, an equivalent way to estimate the PCs would be to conduct a singular value decomposition on a mean centered *G* (McVean 2009). We used the kinship matrix *K* to test for selection on traits using Eq. 8 on the first 22 PCs that, cumulatively, explain 30% of the variation in *K*.

For the denominator of Eq. 8 we used the last half of the PCs (119 to 238). We conducted 200 simulations by simulating traits that evolve neutrally along the kinship matrix using the mvrnorm R function in the MASS package (Venables and Ripley 2002) and testing for selection using Eq. 8.

### GWAS in maize inbreds

We used GEMMA (Zhou and Stephens 2012) to conduct GWAS for trait blups in the GWAS panel, controlling for population structure with a standardized kinship matrix generated by GEMMA (note that this matrix is different from the one used in tests of selection with *Q_PC_*). We conducted two separate GWAS for use in testing for selection in the Ames panel and in the European panel separately. First, for finding SNP associations that we could use to construct breeding values in the Ames panel, we used GBS data for 281 lines from the GWAS panel that had been genotyped by Romay *et al.* (2013). Next, for finding SNP associations for constructing breeding values in the European landraces, we took whole genome data from Bukowski *et al.* (2017) for 263 individuals that had genotype calls for >70% of polymorphic sites and extracted genotypes for sites that overlapped with those present in the European landrace dataset from Unterseer *et al.* (2016). All genotypic data was aligned to v3 of the maize reference genome, except the genotypes of the European landraces, which we lifted over from v2 to v3 using CrossMap (Zhao *et al.* 2013). For both sets of GWAS analyses, tested all SNPs with a minor allele frequency above 0.01, less than 0.05 missing data, and we picked all hits with a likelihood-ratio test p value below 0.005. We pruned both sets of SNPs by using a linkage map from Ogut *et al.* (2015) to construct one cM windows with GenomicRanges in R (Lawrence *et al.* 2013). We picked the SNP with the lowest p value per window and, when multiple SNPs had the same p value, we sampled one SNP randomly.

### Simulated populations and traits

We generated data for Fig. 1 using ms to do coalescent simulations (Hudson 2002) for three large populations of 60 individuals, each with two smaller subpopulations of 30 individuals. Simulations were conducted with the following command: ms 180 1 -t 500 -r 500 10000 -I 6 30 30 30 30 30 30 -ej 0.06 1 2 -ej 0.06 3 4 -ej 0.06 5 6 -ej 0.1 4 6 -ej 0.15 2 6. This command yielded 6,689 loci polymorphic loci for 180 individuals. Neutrally evolving trait values were determined by summing up the number of non-reference allele copies each individual has multiplied by these effect sizes for 50 randomly chosen causal loci (as in Eq. 9. The other 6,639 loci were used to generate a kinship matrix following Eq. 16 and the eigendecomposition of this matrix provided principal components.

### Applying Q_PC_to polygenic scores from the Ames panel and European landraces

We generated combined genetic matrices for each genotyping panel (either Ames or European) and the GWAS panel. In both datasets, we removed sites with a minor allele frequency below 0.01 and a proportion of missing data more than than 0.05 across the combined dataset, leaving 108,110 SNPs in the Ames-GWAS dataset and 441,986 SNPs in the European-GWAS dataset. Missing data points were imputed by replacing each missing genotype from a random sample of the pool of genotypes present in the individuals without missing data. The random imputation step was done once for each missing data point and the same randomly-imputed dataset was used for all subsequent analyses.

We constructed kinship matrices for the Ames panel combined with the GWAS panel and the European panel combined with the GWAS panel following the procedure described in Eq. 16, using 50,000 randomly sampled SNPs with a minor allele frequency > 0.01 and a proportion of missing data below 0.05. The genotype information from the combined datasets was used to construct polygenic scores following Eq. 9.

We used these polygenic scores to test for diversifying selection on 22 traits (described above) for the PCs that cumulatively explained the first 30% of variation in the conditional kinship matrix (182 for the Ames panel and 17 for European maize). As in the *Q_PC_* test, we chose the later 50% of PCs for the denominator. We used Qvalue (Storey *et al.* 2015) to generate false discovery rate estimates (’q values’).

### Simulations of Q_PC_ on polygenic scores

We conducted neutral simulations to detect the rate of false-positive inferences of selection on neutrally-evolving traits. We conducted 800 total: 400 with the Ames panel and 400 with the European landrace panel, and within each panel, 200 with 500 causal loci and 200 with 50 causal loci. For each simulation we simulated a phenotype by randomly picking 500 or 50 sites in the combined genotype datasets and assigning each alternate allele an effect size drawn from a normal distribution with mean 0 and variance 1. For each individual in the GWAS panel, we then calculated a simulated breeding value following Eq. 9. These simulated traits were mapped using a GWAS with the same procedure described above. These simulations showed the GWAS in our sample had relatively low power to detect causal alleles, especially ones of small effect, and also identified a large number of false-positives, as could be expected for a GWAS of this size (Fig. S5).

The loci identified in these GWAS (p < 0.005) were pruned for LD and then used for analysis. We tested for evidence of diversifying selection on polygenic scores in two ways for each set of simulations. First, we used *Q_PC_* with the standard kinship matrix generated using lines in the genotyping set (either Ames or European landraces). Second, we used the conditional *Q_PC_* test described in Eq 11.

We also conducted power simulations using the European landraces. We first simulated traits evolving under diversifying selection by taking trait-associated loci from the neutral simulations and shifting the allele frequencies at these loci in the European landraces based on their latitude of origin. Let *p* be the intial allele frequency in the *j^th^* landrace population, *L* the latitude of the *j^th^* landrace population, *β* the effect size of a alternate allele at the *i^th^* locus, ϵ the mean allele frequency of the *ith* allele, *α* the selection gradient, and *p′* the allele frequency after selection. Then

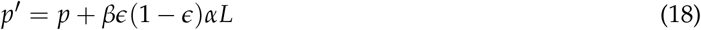

We conducted simulations for three values of *α*: 0.05, 0.01, and 0.005 and tested for selection with conditional *Q_PC_*.

## Data and code availability

All code and data is available at https://github.com/emjosephs/qpc-maize.

## Acknowledgements

We thank Kate Crosby, Cinta Romay, and Peter Bradbury for assistance with maize data, Nancy Chen, Wenbin Mei, and Michelle Stitzer for comments on this manuscript, and members of the Coop, Ross-Ibarra, and Schmitt labs for helpful discussions. E.B.J. is supported by a NSF National Plant Genome Initiative Postdoctoral Research Fellowship (NSF1523733). J.J.B is supported by an NIH F32 NRSA Postdoctoral Research Fellowship (GM126787). J.R.-I. acknowledges support from a NSF PGRP 1238014 and the USDA Hatch project CA-D-PLS-2066-H. G.C. acknowledges support from NIH R01-GM108779, NSF 1262327, and NSF 1353380.

## Appendix 1

### Environmental variation and inferences of selection from *Q_PC_*

Eq. 1 assumes that there is no environmental variation contributing to fitness. It can be rewritten to include the effects of environmental variation as follows:

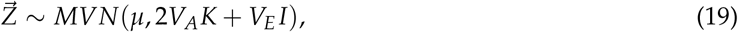

Where 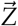 is a vector of traits with mean *μ*, *I* is the identity matrix, *K* is a kinship matrix, and *V_E_* is a constant that measures environmental variation (Falconer and Mackay 1996; Hill 2010). Increases in *V_E_* will thus increase the diagonal entries of the variance-covariance matrix for the multivariate normal distribution of 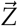. The intuition behind the effect of *V_E_* on the 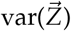 is that, in a properly designed common garden experiment, *V_E_* will increase individual deviations from the expected trait value by increasing the diagonals of the variance-covariance matrix, but will not affect covariance between individuals.

Now, when we mean center 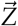 and project it onto a matrix of the eigenvectors of the kinship matrix (*U*), we can get an expression for the set of projections 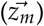 across all eigenvectors:

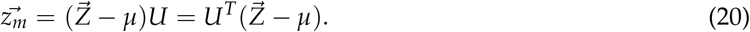

We can express the variance and distribution of *X* as

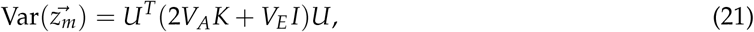

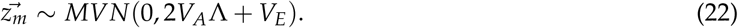

We can standardize *z_m_* by the variance explained by each principal component (the eigenvalues of *K*):

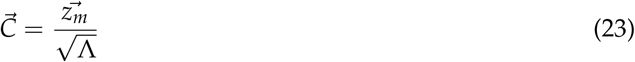

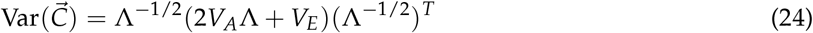

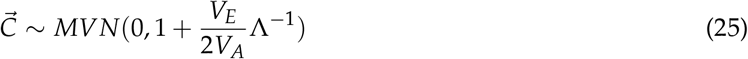

This result suggests that the contribution of *V_E_* will be strongest along PCs with smaller eigenvalues (’later PCs’), so *Q_PC_* is conservative in the face of *V_E_* since it looks for an excess of differentiation along early PCs with larger eigenvalues compared to PCs with smaller eigenvalues.

We tested the intuition described above with simulations of traits that evolve neutrally for with *V_A_* = 1 and *V_E_* = 0, 0.1, and 0.5. We found that increasing *V_E_* increased the variance of *C_M_* at later PCs more than at early PCs (Fig. S1A) and that this meant that fewer simulations showed significant signals of selection than would be expected under neutrality (Fig. S1B)

## Appendix 2

### Additive-by-additive epistasis and *Q_PC_*

We denote the variance contributed by additive-by-additive epistasis as *V_AA_*. Assuming no linkage disequilibrium, we can rewrite Eq. 1 as follows:

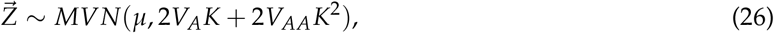

following e.g. Eq. 9.13 in Falconer and Mackay (1996) and Hill (2010). Using the eigendecomposition of *K*, *K* = *U*Λ*U^−^*^1^, where *U* is a matrix whose columns are the eigenvectors of *K* and Λ is a diagonal matrix with the eigenvalues of *K*, we find that *K*^2^ = *U*Λ^2^*U^−^*^1^. As in Appendix 1, we can calculate the 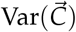 where 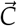 is a vector of the projections of 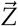 onto *U*, standardized by dividing by Λ^*−*1/2^.

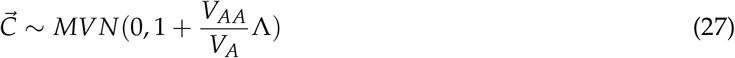

Intuitively, we can see that when *V_AA_* is much larger than *V_A_*, additive-by-additive epistasis will contribute disproportionately to variation along PCs that correspond to higher eigenvalues. Therefore, additive-by-additive epistasis that exceeds *V_A_* can contribute to false positive signals of diversifying selection by increasing trait divergence along earlier PCs. However, in most situations, *V_AA_* is unlikely to be large enough to significantly impact trait variance (Falconer and Mackay 1996; Hill 2010)

## Appendix 3

### Mean centering

Properly mean-centering conditional expectations for polygenic scores and the kinship matrix used to calculate *Q_PC_* on these scores is crucial. However, the choice of how to properly mean-center these two parameters is not entirely straightforward when working with conditional distributions (as in Eq. 10).

To illustrate the problem, imagine that we mean center the conditional expectations for polygenic scores in the genotyping panel 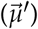 around the mean of the GWAS panel, such that 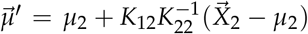 where *μ*_2_ is the mean polygenic score of individuals in the GWAS panel, 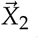 is the vector of polygenic scores in the GWAS panel, and *K*_12_ and *K*_22_ are subsets of the relatedness matrix between individuals in the genotyping panel and GWAS panel as defined for Eq. 10. At the same time, we generate *K*_11_, *K*_22_, and *K*_12_ from the kinship matrix *K* following Eq. 16, where *K* is mean centered around the combined mean of the genotyping and GWAS panel. While these two choices, made separately, seem intuitive, together they lead to a situation where, if *μ*_2_ ≠ *μ*, we can infer signals of adaptive divergence even if none exist. Therefore, we choose to mean center both *K* and 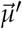 around the mean of all individuals in the genotyping and GWAS panels.

---

**Figure S1.**
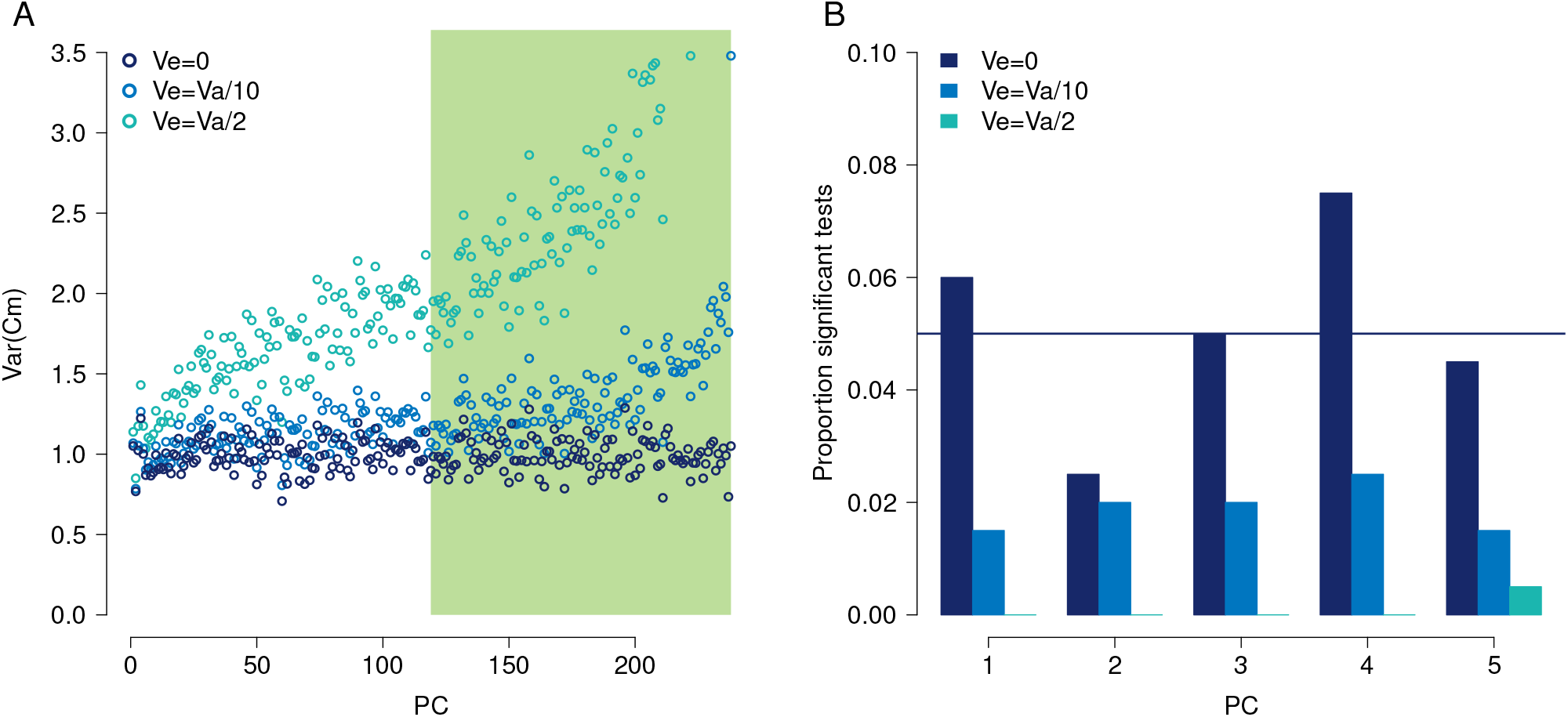
*Q_PC_* on simulated neutral traits with varying amounts of *V_E_*. A) var(*C_m_* across 200 neutral simulations for varying levels of *V_E_*. The PCs used to estimate *V_A_* within populations (the denominator of *Q_PC_*) are shaded green. B) The proportion of 200 neutral simulations that showed evidence of diversifying selection at p < 0.05. We expect that, under neutrality, 0.05 of all simulations should appear significant, but we see that as simulated *V_E_* increases, fewer simulations are significant

**Figure S2.**
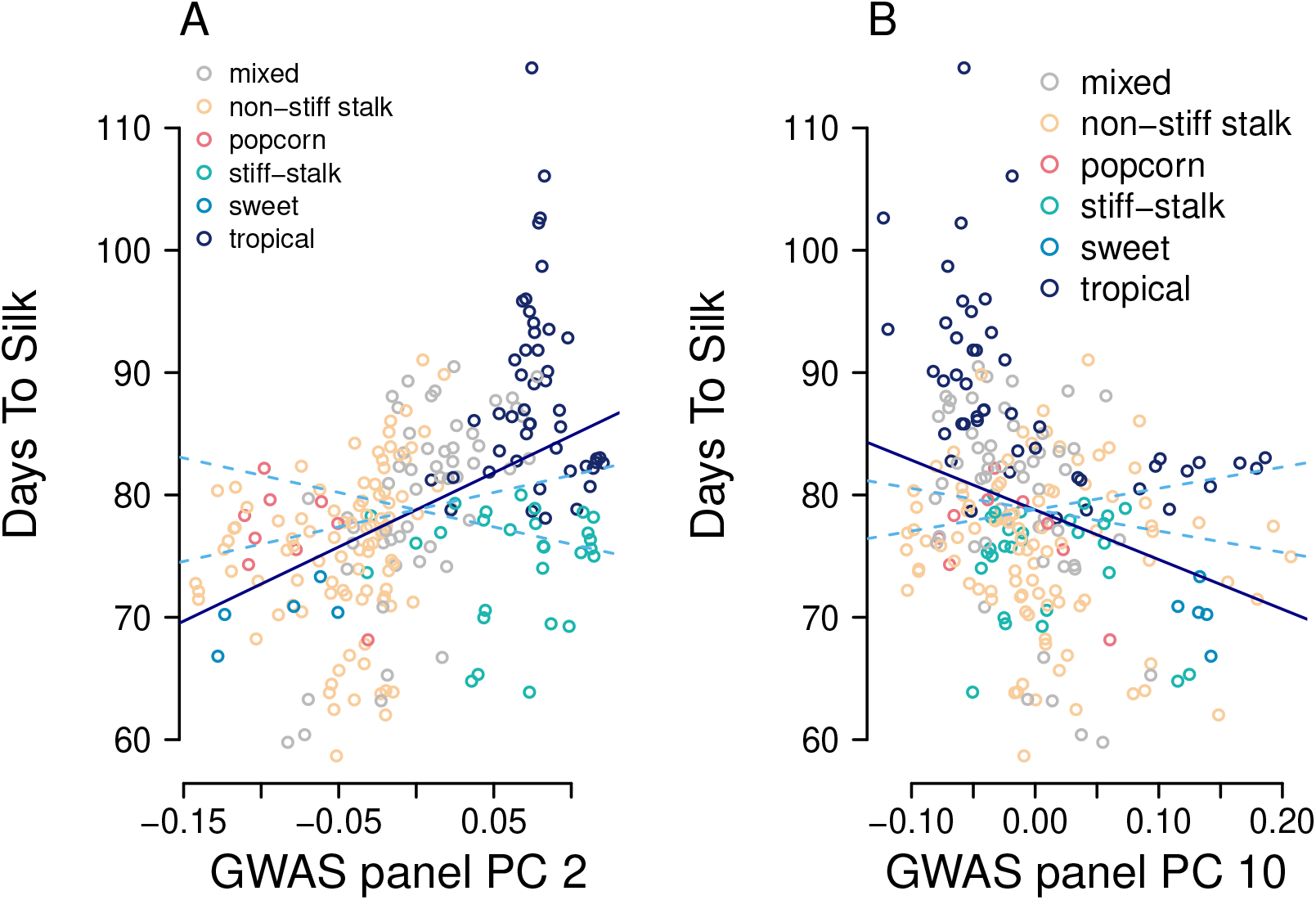
Selection on days to silk along PC10 in the GWAS panel. Each point represents a line in the GWAS panel, colored by its membership in a subpopulation (same colors as Fig. 2A). A) The solid line shows the linear regression of the trait on PC 2 and the dashed lines show the 95% confidence interval of linear regressions expected under neutrality. Note that the linear regression is not the same as the F test done in *Q_PC_*, and that we plot these lines for visualization purposes only. B) The same as (A) but for PC 10.

**Figure S3.**
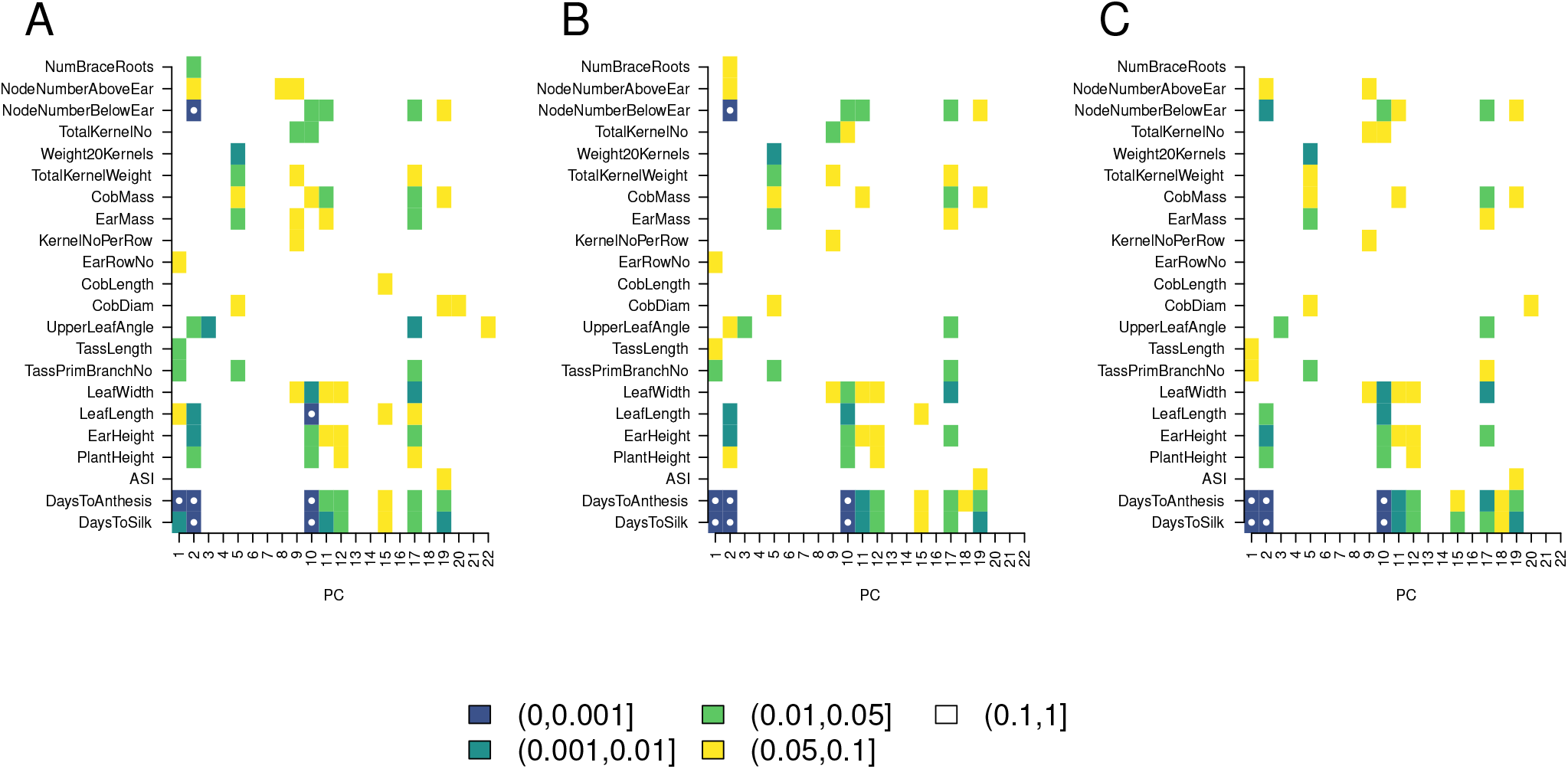
How the PCs used to estimate *V_A_* affect results of *Q_PC_*. We altered the value of *R* from Eq. 8 which changes the number of PCs essentially used to estimate *V_A_* and tested for selection with *Q_PC_*. A) *Q_PC_* results for *R* = 23, so essentially all the PCs past 22 are used in the denominator to estimate *V_A_*. B) The same as A but with *R* 119, so the later half of PCs are used to test for selection. These is the same as the results presented in Fig. 3. C) The same as A and B, but with *R* = 189, so only the latest 50 PCs are used to estimate *V_A_*. Tgit

**Figure S4.**
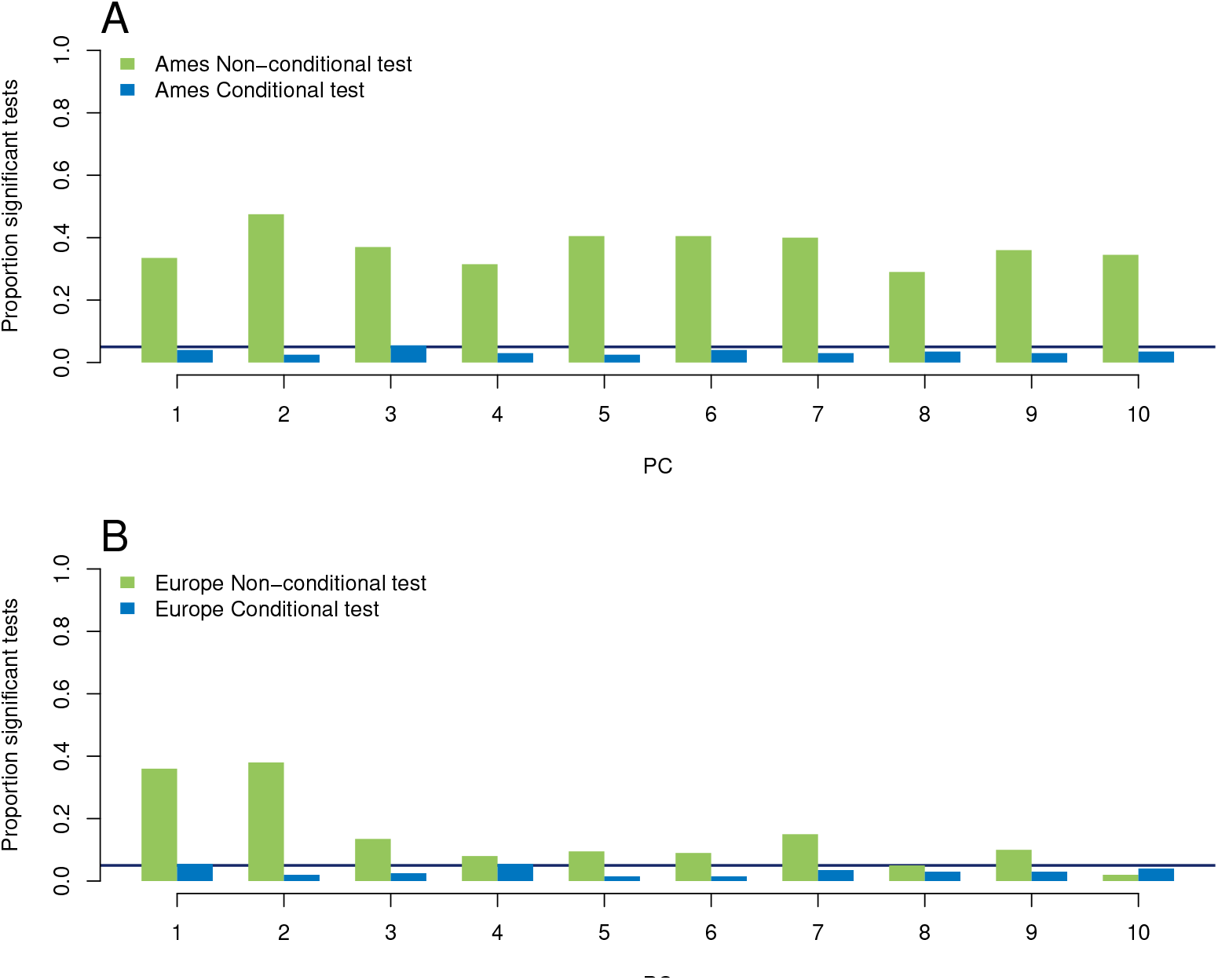
Simulations of *Q_PC_* on polygenic scores where 50 SNPs determine phenotype. A) The proportion of 200 neutral simulations that were significant at the *p* < 0.05 level for the non-conditional *Q_PC_* test and the conditional *Q_PC_* test. A horizontal line is plotted at 0.05, to show the proportion of significant tests expected under the null hypothesis. B) The same information, this time for the European landraces.

**Figure S5.**
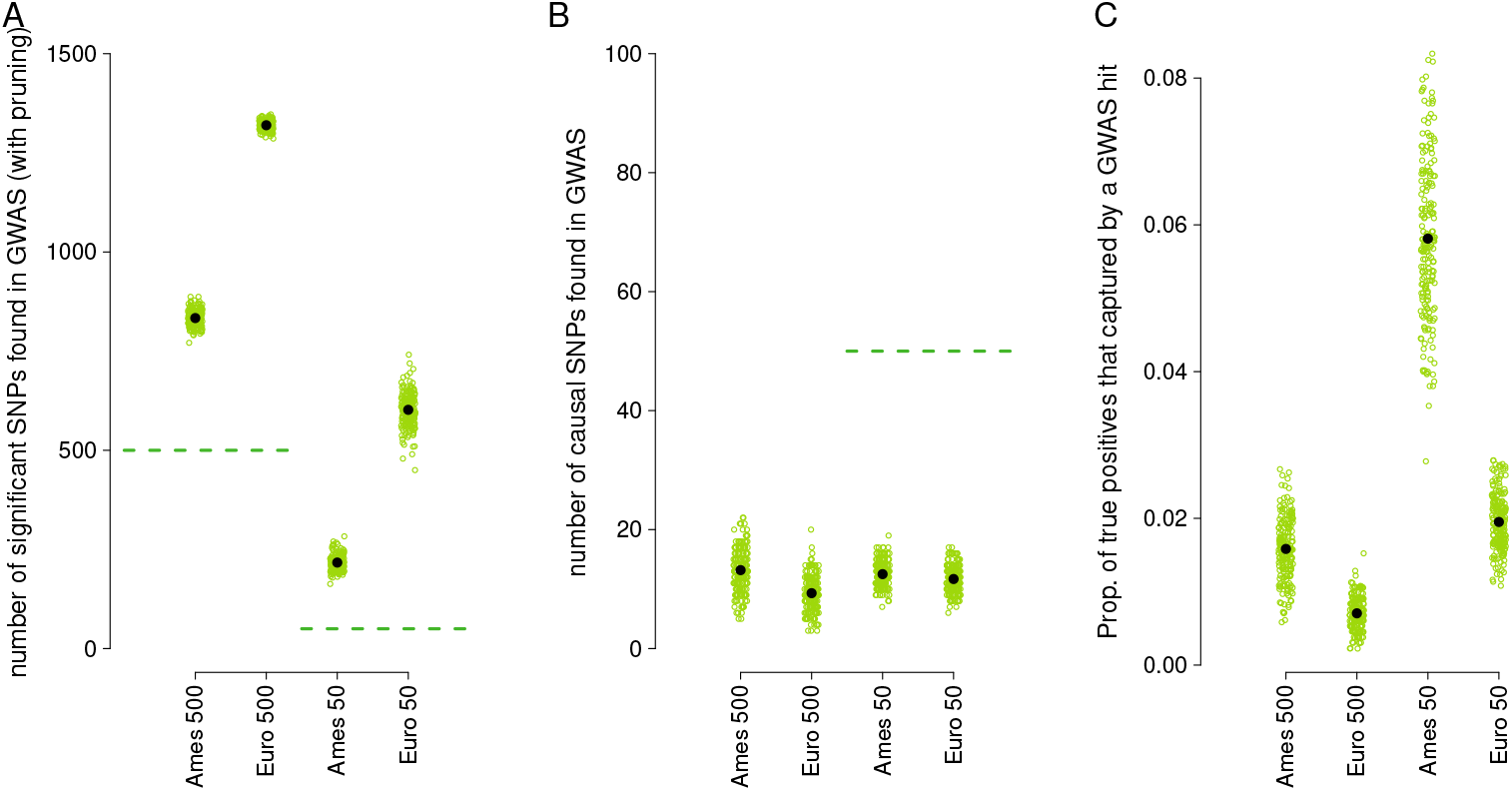
Accuracy of GWAS on simulated traits. A) The number of significant loci that were detected in GWAS (p < 0.005) in each of 200 simulations done using SNPs shared with the Ames panel and shared with the European panel done with 500 or 50 causal loci. In this and subsequent plots, each green dot represents one simulation, the black dots represent the mean across all simulations, and dotted lines represent the expectation if GWAS perfectly found all causal loci with no false positives. B) The number of causal loci from the simulations that were identified in the GWAS at p < 0.005. C) The proportion of simulated causal loci that were in an LD window identified by the GWAS.

**Figure S6.**
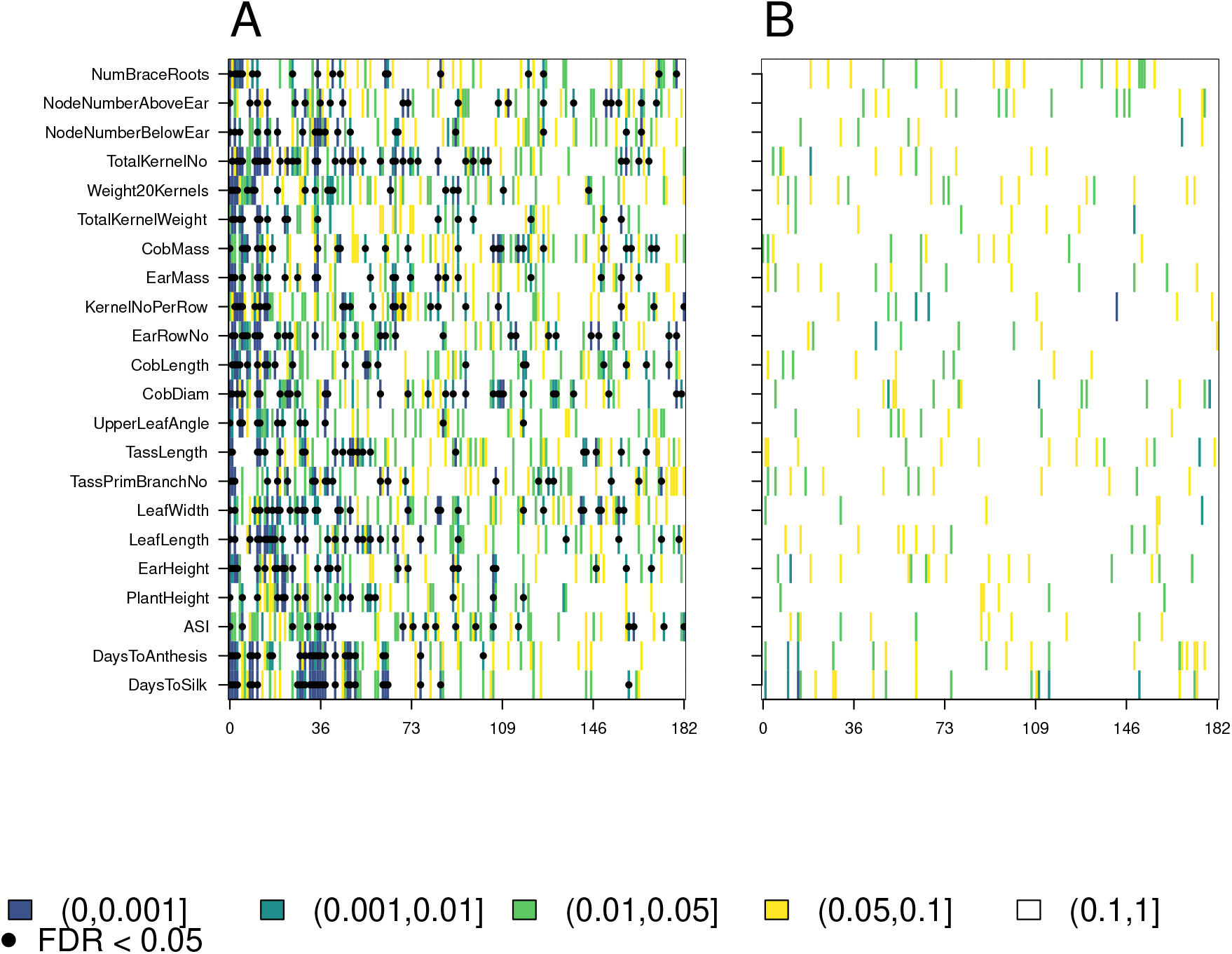
P values from applying *Q_PC_* to polygenic scores in the Ames panel. A) Results from the non-conditional test B) Results from the conditional test

**Figure S7.**
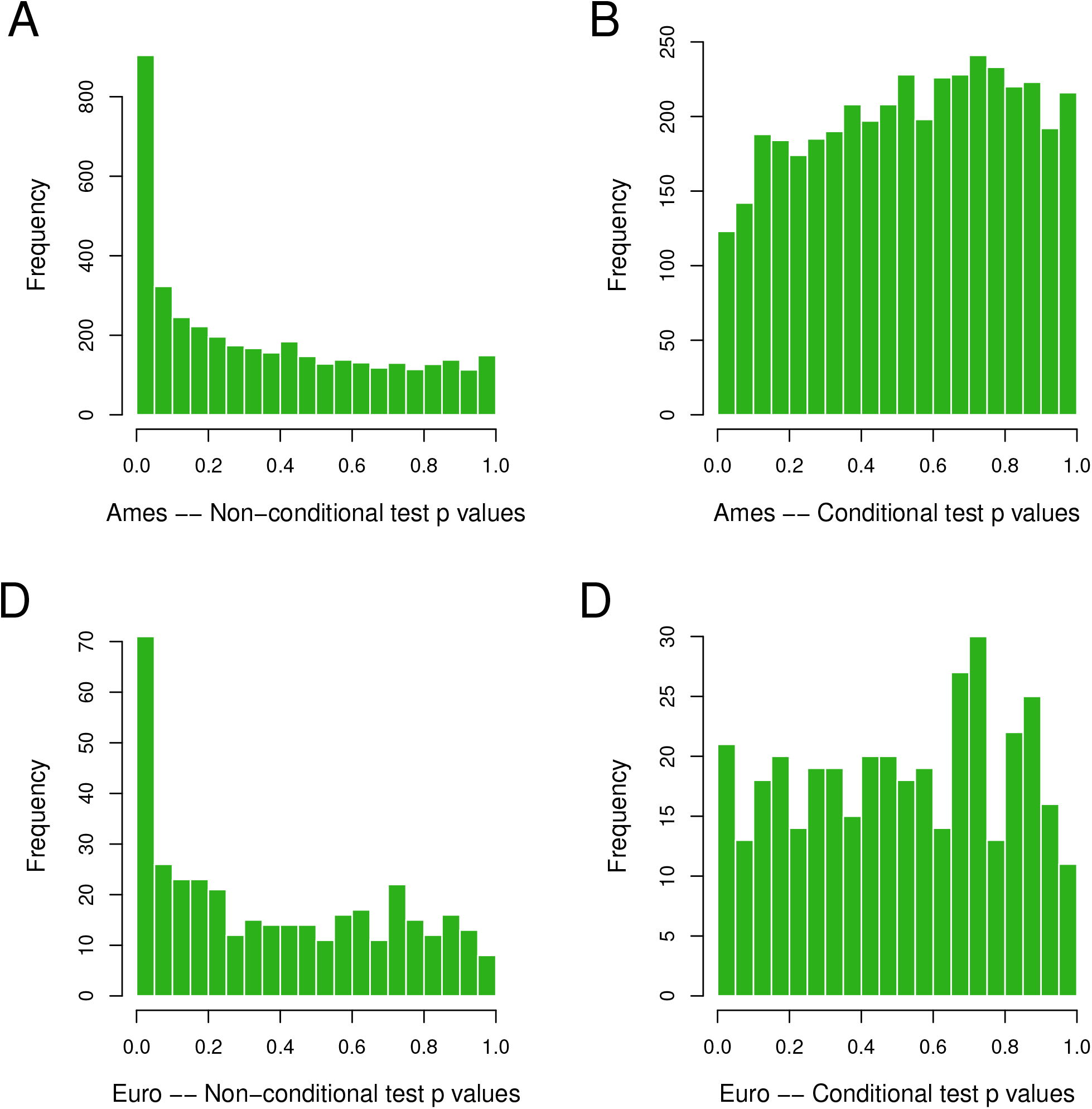
Histograms of P values from applying *Q_PC_* to polygenic scores. A) P values for the non-conditional test in the Ames panel B) P values for the conditional test in the Ames panel. C) P values for the non-conditional test in the European landrace panel. D) P values for the conditional test in the European landrace panel.

## References

Aitken, S. N., S. Yeaman, J. A. Holliday, T. Wang, and S. Curtis-McLane, 2008 Adaptation, migration or extirpation: climate change outcomes for tree populations. Evolutionary Applications 1: 95–111.

Atwell, S., Y. S. Huang, B. J. Vilhjálmsson, G. Willems, M. Horton, et al., 2010 Genome-wide association study of 107 phenotypes in *Arabidopsis thaliana* inbred lines. Nature 465: 627.

Bay, R. A., N. Rose, R. Barrett, L. Bernatchez, C. K. Ghalambor, et al., 2017 Predicting responses to contemporary environmental change using evolutionary response architectures. The American Naturalist 189: 463–473.

Beissinger, T., J. Kruppa, D. Cavero, N.-T. Ha, M. Erbe, et al., 2018 A simple test identifies selection on complex traits. Genetics 209: 321–333.

Berg, J., X. Zhang, and G. Coop, 2017 Polygenic adaptation has impacted multiple anthropometric traits. biorxiv, 167551. Biorxiv.

Berg, J. J. and G. Coop, 2014 A population genetic signal of polygenic adaptation. PLoS Genet. 10: e1004412.

Berg, J. J., A. Harpak, N. Sinnott-Armstrong, A. M. Joergensen, H. Mostafavi, et al., 2018 Reduced signal for polygenic adaptation of height in UK Biobank. bioRxiv.

Brommer, J., 2011 Whither Pst? the approximation of Qst by Pst in evolutionary and conservation biology. Journal of Evolutionary Biology 24: 1160–1168.

Bryc, K., W. Bryc, and J. W. Silverstein, 2013 Separation of the largest eigenvalues in eigenanalysis of genotype data from discrete subpopulations. Theoretical population biology 89: 34–43.

Buckler, E. S., J. B. Holland, P. J. Bradbury, C. B. Acharya, P. J. Brown, et al., 2009 The genetic architecture of maize flowering time. Science 325: 714–718.

Bukowski, R., X. Guo, Y. Lu, C. Zou, B. He, et al., 2017 Construction of the third-generation *Zea mays* haplotype map. GigaScience 7: gix134.

Chen, G.-B., S. H. Lee, Z.-X. Zhu, B. Benyamin, and M. R. Robinson, 2016 EigenGWAS: finding loci under selection through genome-wide association studies of eigenvectors in structured populations. Heredity 117: 51.

Duforet-Frebourg, N., K. Luu, G. Laval, E. Bazin, and M. G. Blum, 2015 Detecting genomic signatures of natural selection with principal component analysis: application to the 1000 genomes data. Molecular Biology and Evolution 33: 1082–1093.

Duvick, D., 2005 Genetic progress in yield of United States maize (*Zea mays L.*). Maydica 50: 193.

Falconer, D. and T. Mackay, 1996 Introduction to Quantitative Genetics. Essex: Benjamin Cummings.

Field, Y., E. A. Boyle, N. Telis, Z. Gao, K. J. Gaulton, et al., 2016 Detection of human adaptation during the past 2000 years. Science p. aag0776.

Flint-Garcia, S. A., A.-C. Thuillet, J. Yu, G. Pressoir, S. M. Romero, et al., 2005 Maize association population: a high-resolution platform for quantitative trait locus dissection. The Plant Journal 44: 1054–1064.

Galinsky, K. J., G. Bhatia, P.-R. Loh, S. Georgiev, S. Mukherjee, et al., 2016 Fast Principal-Component Analysis Reveals Convergent Evolution of ADH1B in Europe and East Asia. American Journal of Human Genetics 98: 456–472.

Hadfield, J. and S. Nakagawa, 2010 General quantitative genetic methods for comparative biology: phylogenies, taxonomies and multi-trait models for continuous and categorical characters. Journal of Evolutionary Biology 23: 494–508.

Henderson, C. R., 1950 Estimation of genetic parameters. In Biometrics, volume 6, pp. 186–187, International Biometric Soc.

Henderson, C. R., 1953 Estimation of variance and covariance components. Biometrics 9: 226–252.

Hereford, J., 2009 A quantitative survey of local adaptation and fitness trade-offs. The American Naturalist 173: 579–588.

Hill, W. G., 2010 Understanding and using quantitative genetic variation. Philosophical Transactions of the Royal Society of London B: Biological Sciences 365: 73–85.

Hill, W. G., M. E. Goddard, and P. M. Visscher, 2008 Data and theory point to mainly additive genetic variance for complex traits. PLoS genetics 4: e1000008.

Howden, S. M., J.-F. Soussana, F. N. Tubiello, N. Chhetri, M. Dunlop, et al., 2007 Adapting agriculture to climate change. Proceedings of the National Academy of Sciences 104: 19691–19696.

Hudson, R. R., 2002 Generating samples under a wright–fisher neutral model of genetic variation. Bioinformatics 18: 337–338.

Hung, H., C. Browne, K. Guill, N. Coles, M. Eller, et al., 2012 The relationship between parental genetic or phenotypic divergence and progeny variation in the maize nested association mapping population. Heredity 108: 490.

Karhunen, M., J. Merilä, T. Leinonen, J. Cano, and O. Ovaskainen, 2013 DRIFTSEL: an R package for detecting signals of natural selection in quantitative traits. Molecular Ecology Resources 13: 746–754.

Kremer, A. and V. Le Corre, 2012 Decoupling of differentiation between traits and their underlying genes in response to divergent selection. Heredity 108: 375.

Latta, R. G., 1998 Differentiation of allelic frequencies at quantitative trait loci affecting locally adaptive traits. The American Naturalist 151: 283–292.

Lawrence, M., W. Huber, H. Pages, P. Aboyoun, M. Carlson, et al., 2013 Software for computing and annotating genomic ranges. PLoS Computational Biology 9: e1003118.

Le Corre, V. and A. Kremer, 2012 The genetic differentiation at quantitative trait loci under local adaptation. Molecular Ecology 21: 1548–1566.

Leimu, R. and M. Fischer, 2008 A meta-analysis of local adaptation in plants. PloS one 3: e4010.

Leinonen, T., R. S. McCairns, R. B. O’hara, and J. Merilä, 2013 Qst–Fst comparisons: evolutionary and ecological insights from genomic heterogeneity. Nature Reviews Genetics 14: 179.

Leinonen, T., R. B. Ohara, J. Cano, and J. Merilä, 2008 Comparative studies of quantitative trait and neutral marker divergence: a meta-analysis. Journal of Evolutionary Biology 21: 1–17.

Liu, X., P.-R. Loh, L. J. O’Connor, S. Gazal, A. Schoech, et al., 2018 Quantification of genetic components of population differentiation in UK Biobank traits reveals signals of polygenic selection. bioRxiv.

Luu, K., E. Bazin, and M. G. Blum, 2016 pcadapt: an R package to perform genome scans for selection based on principal component analysis. Molecular Ecology Resources 17: 67–77.

McVean, G., 2009 A genealogical interpretation of principal components analysis. PLoS Genetics 5: e1000686.

Menozzi, P., A. Piazza, and L. Cavalli-Sforza, 1978 Synthetic maps of human gene frequencies in Europeans. Science 201: 786–792.

Mikel, M. A. and J. W. Dudley, 2006 Evolution of North American dent corn from public to proprietary germplasm. Crop Science 46: 1193–1205.

Mitchell-Olds, T., J. H. Willis, and D. B. Goldstein, 2007 Which evolutionary processes influence natural genetic variation for phenotypic traits? Nature Reviews Genetics 8: 845.

Novembre, J. and N. H. Barton, 2018 Tread lightly interpreting polygenic tests of selection. Genetics 208: 1351–1355.

Novembre, J., T. Johnson, K. Bryc, Z. Kutalik, A. R. Boyko, et al., 2008 Genes mirror geography within Europe. Nature 456: 98.

Novembre, J. and M. Stephens, 2008 Interpreting principal component analyses of spatial population genetic variation. Nature Genetics 40: 646–649.

Ogut, F., Y. Bian, P. J. Bradbury, and J. B. Holland, 2015 Joint-multiple family linkage analysis predicts within-family variation better than single-family analysis of the maize nested association mapping population. Heredity 114: 552.

Ovaskainen, O., M. Karhunen, C. Zheng, J. M. C. Arias, and J. Merilä, 2011 A new method to uncover signatures of divergent and stabilizing selection in quantitative traits. Genetics 189: 621–632.

Patterson, N., A. L. Price, and D. Reich, 2006 Population structure and eigenanalysis. PLoS Genetics 2: e190.

Peiffer, J. A., M. C. Romay, M. A. Gore, S. A. Flint-Garcia, Z. Zhang, et al., 2014 The genetic architecture of maize height. Genetics 196: 1337–1356.

Price, A. L., N. J. Patterson, R. M. Plenge, M. E. Weinblatt, N. A. Shadick, et al., 2006 Principal components analysis corrects for stratification in genome-wide association studies. Nature genetics 38: 904.

Prout, T. and J. Barker, 1993 F statistics in *Drosophila buzzatii*: selection, population size and inbreeding. Genetics 134: 369–375.

Pujol, B., A. J. Wilson, R. Ross, and J. Pannell, 2008 Are Qst–Fst comparisons for natural populations meaningful? Molecular Ecology 17: 4782–4785.

R Core Team, 2018 R: A Language and Environment for Statistical Computing. R Foundation for Statistical Computing, Vienna, Austria.

Robinson, M. R., G. Hemani, C. Medina-Gomez, M. Mezzavilla, T. Esko, et al., 2015 Population genetic differentiation of height and body mass index across Europe. Nature Genetics 47: 1357.

Romay, M. C., M. J. Millard, J. C. Glaubitz, J. A. Peiffer, K. L. Swarts, et al., 2013 Comprehensive genotyping of the USA national maize inbred seed bank. Genome biology 14: R55.

Savolainen, O., M. Lascoux, and J. Merilä, 2013 Ecological genomics of local adaptation. Nature Reviews Genetics 14: 807.

Sohail, M., R. M. Maier, A. Ganna, A. Bloemendal, A. R. Martin, et al., 2018 Signals of polygenic adaptation on height have been overestimated due to uncorrected population structure in genome-wide association studies. bioRxiv.

Spitze, K., 1993 Population structure in *Daphnia obtusa*: quantitative genetic and allozymic variation. Genetics 135: 367–374.

Storey, J. D., A. J. Bass, A. Dabney, and D. Robinson, 2015 qvalue: Q-value estimation for false discovery rate control. R package version 2.8.0.

Swarts, K., R. M. Gutaker, B. Benz, M. Blake, R. Bukowski, et al., 2017 Genomic estimation of complex traits reveals ancient maize adaptation to temperate North America. Science 357: 512–515.

Takeda, S. and M. Matsuoka, 2008 Genetic approaches to crop improvement: responding to environmental and population changes. Nature Reviews Genetics 9: 444.

Tenaillon, M. I. and A. Charcosset, 2011 A European perspective on maize history. Comptes Rendus Biologies 334: 221–228.

Thompson, R., 2008 Estimation of quantitative genetic parameters. Proceedings of the Royal Society of London B: Biological Sciences 275: 679–686.

Turchin, M. C., C. W. Chiang, C. D. Palmer, S. Sankararaman, D. Reich, et al., 2012 Evidence of widespread selection on standing variation in Europe at height-associated SNPs. Nature Genetics 44: 1015.

Unterseer, S., S. D. Pophaly, R. Peis, P. Westermeier, M. Mayer, et al., 2016 A comprehensive study of the genomic differentiation between temperate dent and flint maize. Genome Biology 17: 137.

Venables, W. N. and B. D. Ripley, 2002 Modern Applied Statistics with S. Springer, New York, fourth edition, ISBN 0-387-95457-0.

Wang, W., R. Mauleon, Z. Hu, D. Chebotarov, S. Tai, et al., 2018 Genomic variation in 3,010 diverse accessions of Asian cultivated rice. Nature 557: 43.

Whitlock, M. C., 2008 Evolutionary inference from Qst. Molecular Ecology 17: 1885–1896.

Whitlock, M. C. and K. J. Gilbert, 2012 Qst in a hierarchically structured population. Molecular Ecology Resources 12: 481–483.

Wickham, H., R. Francois, L. Henry, and K. Müller, 2017 dplyr: A Grammar of Data Manipulation. R package version 0.7.4.

Yu, J., G. Pressoir, W. H. Briggs, I. V. Bi, M. Yamasaki, et al., 2006 A unified mixed-model method for association mapping that accounts for multiple levels of relatedness. Nature Genetics 38: 203.

Zhao, H., Z. Sun, J. Wang, H. Huang, J.-P. Kocher, et al., 2013 Crossmap: a versatile tool for coordinate conversion between genome assemblies. Bioinformatics 30: 1006–1007.

Zhou, X. and M. Stephens, 2012 Genome-wide efficient mixed-model analysis for association studies. Nature Genetics 44: 821.

